# The plastomes of *Hyalomonas oviformis* and *Hyalogonium fusiforme* evolved dissimilar architecture after the loss of photosynthesis

**DOI:** 10.1101/2022.03.29.486296

**Authors:** Alexandra E. DeShaw, Francisco Figueroa-Martinez, Thomas Pröschold, Maike Lorenz, Aurora M. Nedelcu, David R. Smith, Adrián Reyes-Prieto

## Abstract

The loss of photosynthesis in land plants and algae is typically associated with parasitism but can also occur in free-living species, including chlamydomonadalean green algae. The plastid genomes (ptDNAs) of colorless chlamydomonadalean species are surprisingly diverse in architecture, including highly expanded forms (*Polytoma uvella, Leontynka pallida*) as well as outright genome loss (*Polytomella* species). Here, we explore the ptDNAs of *Hyalomonas (Hm.) oviformis* (SAG 62-27; formerly known as *Polytoma oviforme*) and *Hyalogonium (Hg.) fusiforme* (SAG 62-1c), each representing independent losses of photosynthesis within the Chlamydomonadales. The *Hm. oviformis* ptDNA is moderately sized (132 kb), smaller than that of its photosynthetic relative *Hyalomonas chlamydogama* SAG 11-48b (198.3 kb), with a reduced gene complement but still encoding the ATPase subunits. The *Hg. fusiforme* plastome, however, is the largest yet observed in colorless plants or algae (~463 kb) and has a coding repertoire that is almost identical to that of its photosynthetic relatives in the genus *Chlorogonium*. Furthermore, the ptDNA of *Hg. fusiforme* shows no clear evidence of pseudogenization, which is consistent with our analyses showing that *Hg. fusiforme* is the non-photosynthetic lineage of most recent origin among the known colorless Chlamydomonadales. Together, these new ptDNAs clearly show that, in contrast to parasitic algae, plastid genome compaction is not an obligatory route following the loss of photosynthesis in free-living algae, and that certain chlamydomonadalean algae have a remarkable propensity for genomic expansion, which can persist regardless of the trophic strategy.

**One sentence summary:** The plastid genomes of two free-living chlamydomonadalean algae, *Hyalomonas oviformis* and *Hyalogonium fusiforme*, reveal different evolutionary stages following the loss of photosynthesis.

## Introduction

The loss of photosynthesis has occurred many times throughout eukaryotic evolution (Figueroa-Martinez et al., 2015) and is typically associated with a switch to a parasitic or pathogenic lifestyle in both land plants and algae (Wicke et al., 2013). Nevertheless, the transition to a nonphotosynthetic lifestyle has also occurred in free-living species, such as in various members of the green algal order Chlamydomonadales (Chlorophyceae). There are at least six distinct, free-living chlamydomonadalean lineages that have independently lost photosynthetic capabilities, including species in the genera *Polytomella*, *Polytoma*, *Leontynka*, the isolate *Volvocales sp*. NrCl902, *Hyalogonium fusiforme* and *Hylaomonas oviformis* gen. et sp. nov (Nedelcu, 2001; Vernon et al., 2001; Figueroa-Martinez et al., 2017; Kayama et al., 2020, Pánek et al., 2022; Figure 1).

**Figure 1.**
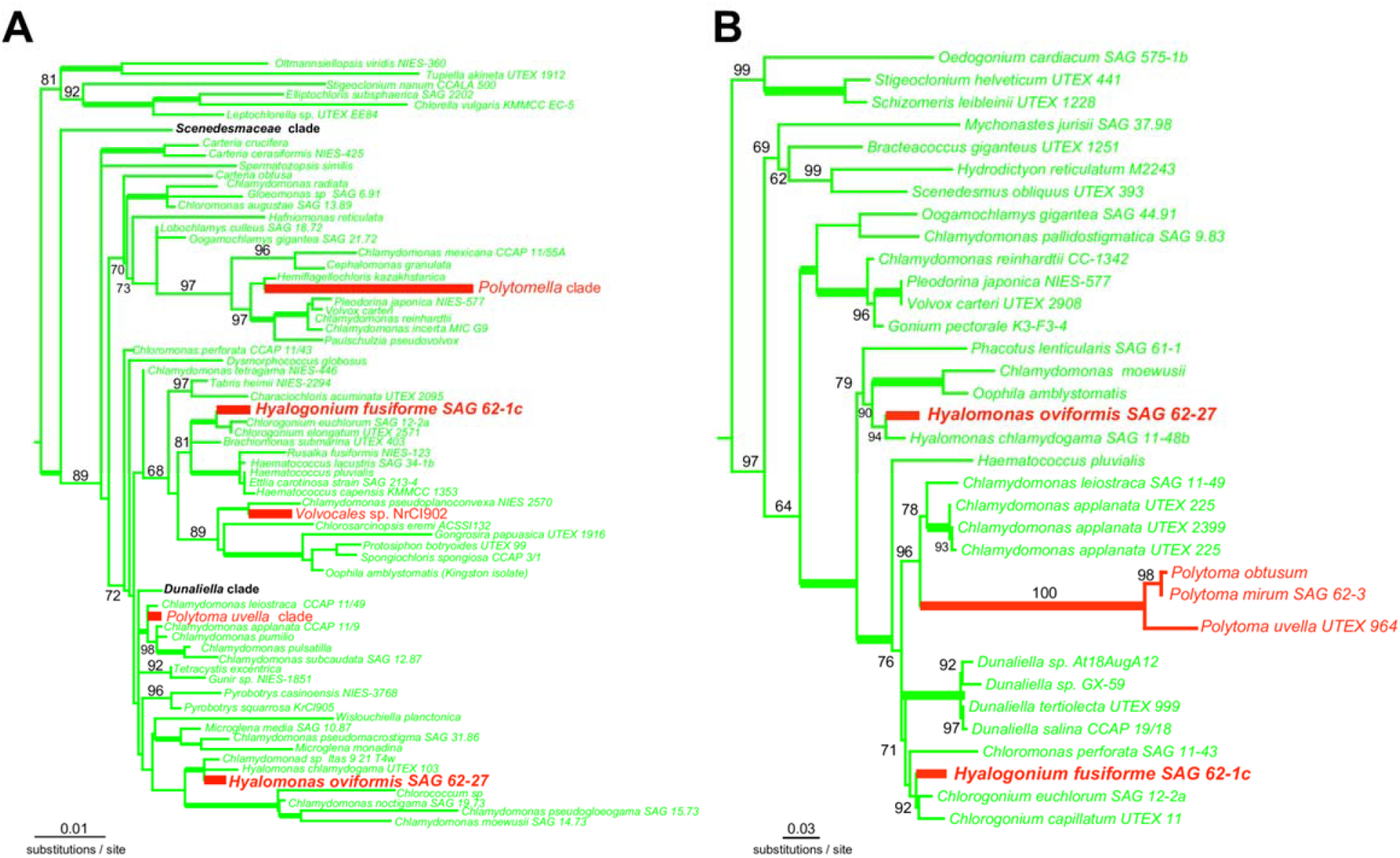
Phylogenetic trees estimated from **A**) nuclear 18S rRNA and **B**) plastid 16S rRNA gene sequences. Both maximum likelihood (ML) trees were estimated with IQ-TREE considering the best fitting nucleotide substitution model in each case (see text for details). The 18S rRNA ML tree was rooted using the Trebouxiophyceae clade as outgroup, in the case of the 16S rRNA tree the outgroup reference were the three members of the OCC (Oedogoniales, Chaetophorales and Chaetopeltidales) clade. Branch lengths are proportional to the number of substitutions per site as indicated by the corresponding scale bars. Numbers near nodes indicate values of bootstrap proportion support (10,000 ultra-fast replicates) and thick branches denote 100% bootstrap. Nonphotosynthetic chlamydomonadalean species are indicated in orange; green branches represent photosynthetic/mixotrophic lineages.

Previous studies of colorless Chlamydomonadales have identified disparate and intriguing patterns of plastid genome (ptDNA) evolution. *Polytomella* species, for instance, retain a colorless plastid but no longer have ptDNA (Smith & Lee, 2014). *Polytoma uvella* on the other hand, has a highly expanded plastome (~240 kb), resulting from an abundance of noncoding DNA (Figueroa-Martinez et al., 2017). In fact, despite having lost all genes related to photosynthesis, this ptDNA is larger than those of its close photosynthetic relative, *Chlamydomonas leiostraca* (Figueroa-Martinez et al., 2017). However, *Polytomella* and *P. uvella* belong to just two of the six known independently evolved non-photosynthetic chlamydomondalean algae (Figure 1). To improve our understanding of the evolutionary mechanisms underlying the changes in ptDNA organization associated with the loss of photosynthesis in free-living species, we explored ptDNA evolution in *Polytoma* sp. SAG 62-27 (hereafter *Hyalomonas oviformis* gen. et sp. nov.; Figure 2A) and *Hyalogonium fusiforme* SAG 62-1c (Figure 2B), which represent two distinct and poorly studied nonphotosynthetic lineages within the Chlamydomonadales. These colorless algae are closely related to the photosynthetic species *Chlamydomonas chlamydogama* (hereafter transferred to *Hyalomonas*) and *Chlorogonium* spp., respectively. We found that the *Hm. oviformis* ptDNA is moderately sized with a reduced gene content typical of other nonphotosynthetic plastomes. The *Hg. fusiforme* ptDNA, however, retains a near-complete collection of genes involved in photosynthesis with no obvious examples of pseudogenization. Even more surprising, it represents one of the largest plastomes yet observed in a nonphotosynthetic plant or algae.

**Figure 2.**
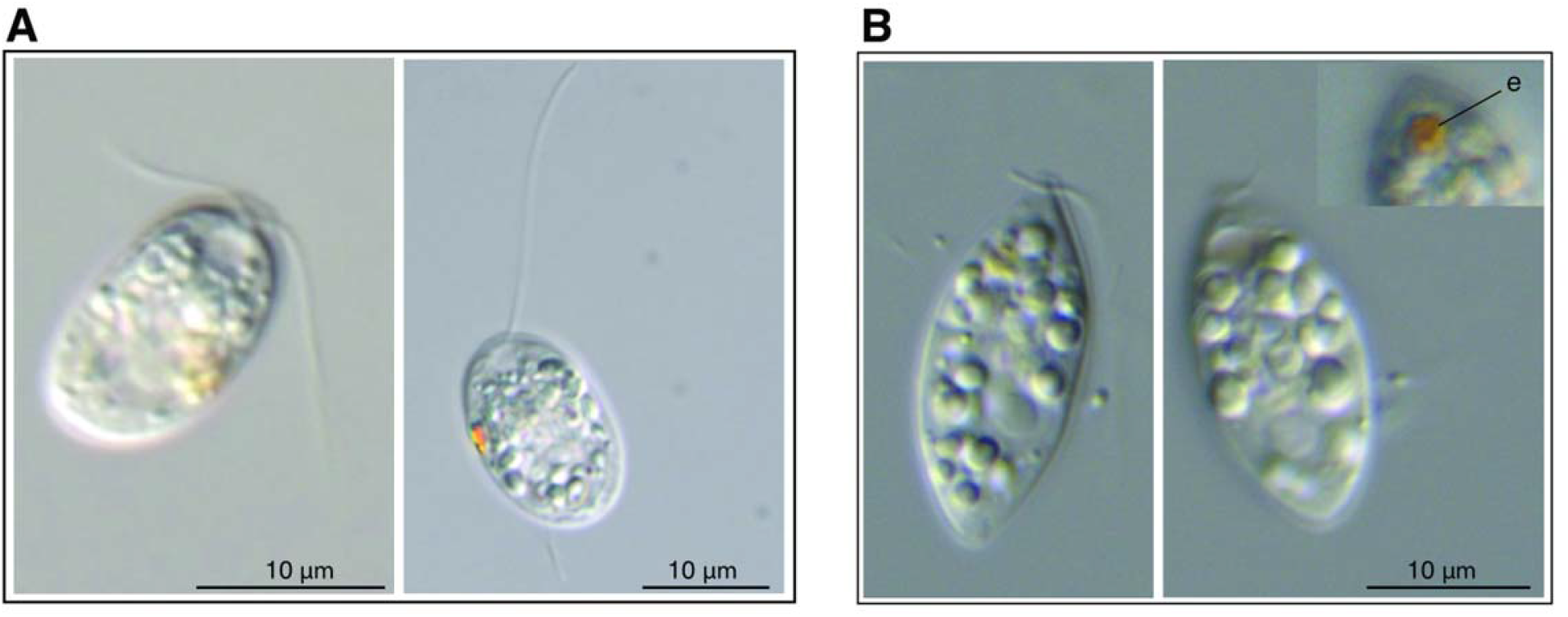
Differential interference contrast micrographs of **A**) *Hyalomonas oviformis* (SAG 62-27) in low-melting point agarose and **B**) *Hyalogonium fusiforme* (SAG 62-1c), with the carotene-rich eyespot (e) indicated in the inset image.

## Results and Discussion

### Hyalomonas, a new non-photosynthetic genus

The strain SAG 62-27 was originally assigned to the genus *Polytoma* with the species epithet *oviforme* Pringsheim (nom. nud., without valid description; Koch 1964). Phylogenetic analyses revealed that this taxon is not closely related to the type species of *Polytoma, P. uvella*, and represents therefore a new genus (Nedelcu, 2001; Figueroa-Martinez et al., 2017). The other species that belongs to this new genus is the photosynthetic species *Chlamydomonas chlamydogama* originally described by Bold 1949.

### *Hyalomonas* gen. nov

Description: Cells biflagellated, with chloroplast of *Agloe*-type containing a single pyrenoid or reduced parietal leucoplast. Cells surrounded with glycoprotein cell wall. Plastids with prominent eyespot. Asexual reproduction by 4-8 zoospores. Sexual reproduction by isogamy.

Diagnosis: Differs from other genera of biflagellated chlorophytes by SSU-ITS and plastome sequence.

Type species (designated here): *Hyalomonas oviformis* sp. nov.

### *Hyalomonas oviformis* sp. nov

Description: Cells oviform, 11-15 μm long, 8-10 μm wide, two flagella with 1-1½ body length, with parietal leucoplast containing spot-like anterior eyespot. Nucleus in posterior position. Two contractile vacuoles (Figure 2A).

Diagnosis: Differs from other species of non-photosynthetic biflagellated chlorophytes by SSU-ITS (GenBank accession OM985702) and plastome (GenBank accession XXXXXX) sequences.

Holotype (designated here): The strain SAG 62-27 cryopreserved in metabolically inactive state at the Culture Collection of Algae (SAG), University Göttingen, Germany.

### *Hyalomonas chlamydogama* (Bold) comb. nov

Basionym: *Chlamydomonas chlamydogama* Bold 1949, *Bull. Torrey Bot. Cl*. **76**, 101, fig. 1 (lectotype designated here).

Epitype (designated here): The strain SAG 11-48b cryopreserved in metabolically inactive state at the Culture Collection of Algae (SAG), University Göttingen, Germany.

### The plastomes of two non-photosynthetic chlamydomonadalean lineages of distinct age

Our phylogenetic analyses show that *Hm. oviformis* and *Hg. fusiforme* lost photosynthesis independently from one another as well as from other colorless lineages within the Chlamydomonadales (Figure 2). Among the closest known photosynthetic relatives of *Hm. oviformis* and *Hg. fusiforme* are *Hm*. *chlamydogama* and *Cg*. *euchlorum*, respectively (Figure 2). *RelTime* analyses with protein and rDNA sequence data suggest that *Hm. oviformis* and *P. uvella* likely diverged from their respective photosynthetic siblings during close, likely overlapping periods (~90-120 MYA) (Figures S1, S4, S5; Table S1). However, our analyses consistently recovered *Hg. fusiforme* as the colorless lineage of most recent origin among the chlamydomonadalean taxa investigated here (~45-75 MYA) (Figure S1).

To understand the consequences of the loss of photosynthesis in these two non-photosynthetic lineages of distinct evolutionary age, we sequenced the ptDNAs of *Hm. oviformis* and *Hg. fusiforme* as well as those of *Hm. chlamydogama* and *Cg. euchlorum*. The plastomes of *Hm. oviformis* (132 kb) and *Hm. chlamydogama* (198.3 kb) were assembled into circular-mapping molecules (Figure 3; Table S2). Unlike *Hm. chlamydogama*, the *Hm. oviformis* ptDNA lacks inverted repeats, which is a recurring theme among plastomes of nonphotosynthetic plants and algae (Figueroa-Martinez et al., 2015, 2017; Wicke et al., 2013). However, the presence of a large number of repeats prevented us from generating intact assemblies of the *Hg. fusiforme* and *Cg. euchlorum* ptDNAs, despite employing long-read PacBio sequencing. The *Hg. fusiforme* and *Cg. euchlorum* plastome sequences are distributed across 43 and 4 scaffolds with accumulative lengths of 463 kb and 314 kb, respectively (Figure S2; Table S2).

**Figure 3.**
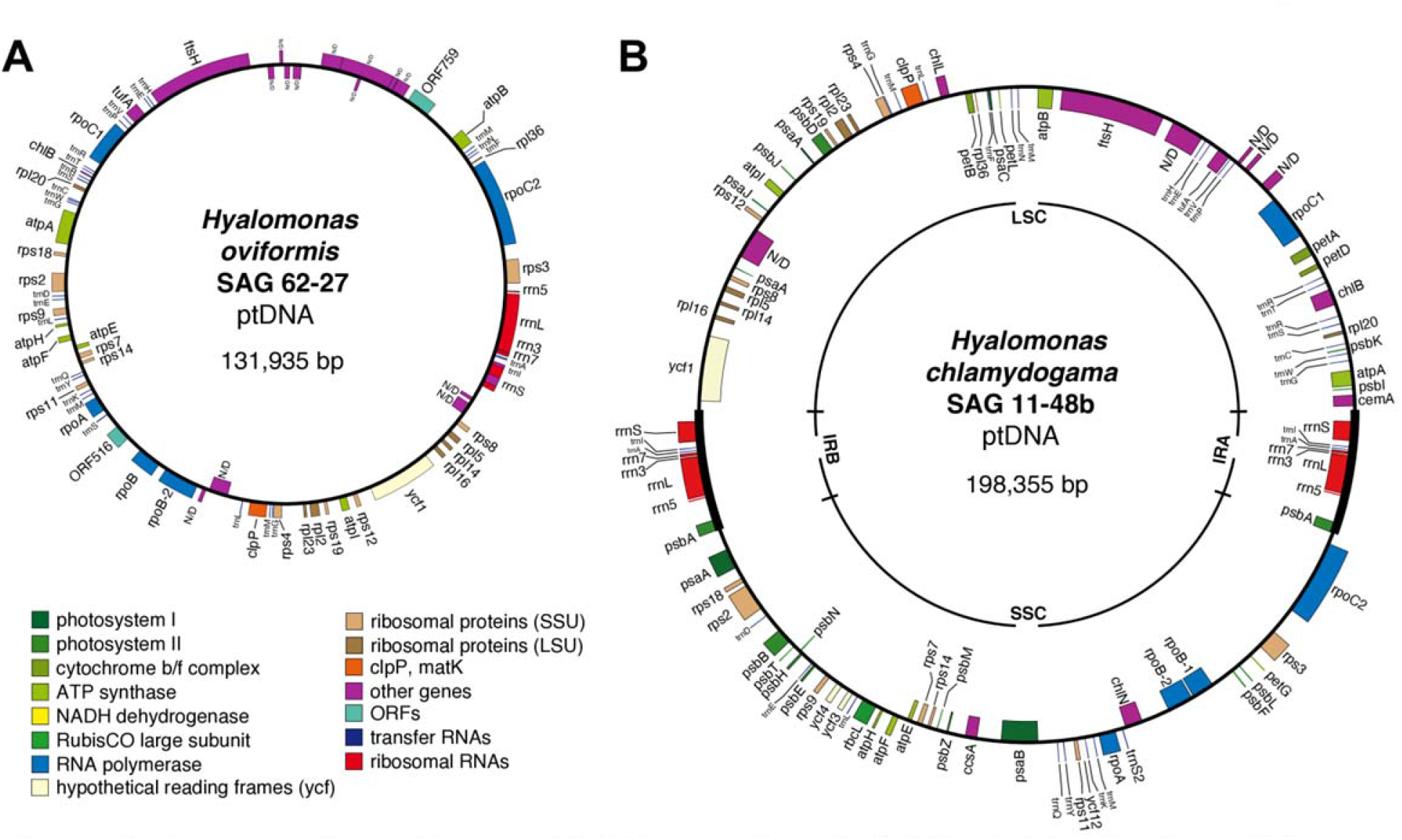
Circular gene maps of the plastid genomes of **A**) *Hyalomonas oviformis* (SAG 62-27) and **B**) *Hm. chlamydogama* (SAG 11-48b). Large single copy (LSC), small single copy (SSC) and inverted repeat (IR*N*) regions are indicated. The graphic representations were prepared with the OrganellarGenomeDRAW (OGDRAW) v1.3.1 tool (Greiner et al., 2019) implemented in the CHLOROBOX server (https://chlorobox.mpimp-golm.mpg.de/index.html).

Chlamydomonadalean ptDNAs are renowned for being extremely hard to assemble (Gaouda et al., 2018), with many reported assemblies existing in highly fragmented forms (Figueroa-Martinez et al., 2017; Gaouda et al., 2018). Our read coverage analyses indicate that all the *Hg. fusiforme* and *Cg. euchlorum* scaffolds are genuine components of the plastid genome and not ptDNA-like sequences from the mitochondrial or nuclear genomes. The read coverage survey also supports the existence of an inverted repeat in *Hg. fusiforme* (Table S3).

### Comparative plastid genomics among photosynthetic and nonphotosynthetic species

To gain a better understanding of the evolutionary mechanisms that shaped the genetic architecture of non-photosynthetic plastomes in Chlamydomonadales, we compared the ptDNAs of the two non-photosynthetic species with those of their closest known photosynthetic relatives as well as other colorless species. The *Hm. oviformis* plastome is circa 65 kb smaller than that of the photosynthetic *Hm. chlamydogama* but still large when compared to the ptDNAs from other colorless plants and algae, which are typically under 75 kb. One of the reasons for this is that *Hm. oviformis* encodes all subunits of the cF_0_-CF_1_ ATPase (*atpA, atpB, atpE, atpF, atpH*, and *atpI*). These genes are similar in length to their orthologues in *Hm. chlamydogama*, but some of them (4/6) show elevated rates of nucleotide substitution (Figure 4). The genes of the *atp* series are not generally retained in colorless plastids, but notable exceptions to varying degrees come from the chlamydomonadalean *Leontynka pallida* (Pánek et al., 2022), other diverse colorless algae (Tanifuji et al., 2020) and angiosperms (Wicke et al., 2013). Nevertheless, like the plastomes of other colorless plants and algae, the *Hm. oviformis* ptDNA has lost all genes encoding subunits of the two photosystems, the *b_6_f* complex, and the enzymes for chlorophyll biosynthesis (Table S4). However, it is still possible to distinguish conserved collinear plastid gene arrangements between *Hm. oviformis* and the photosynthetic *Hm. chlamydogama* (Figure S3).

**Figure 4.**
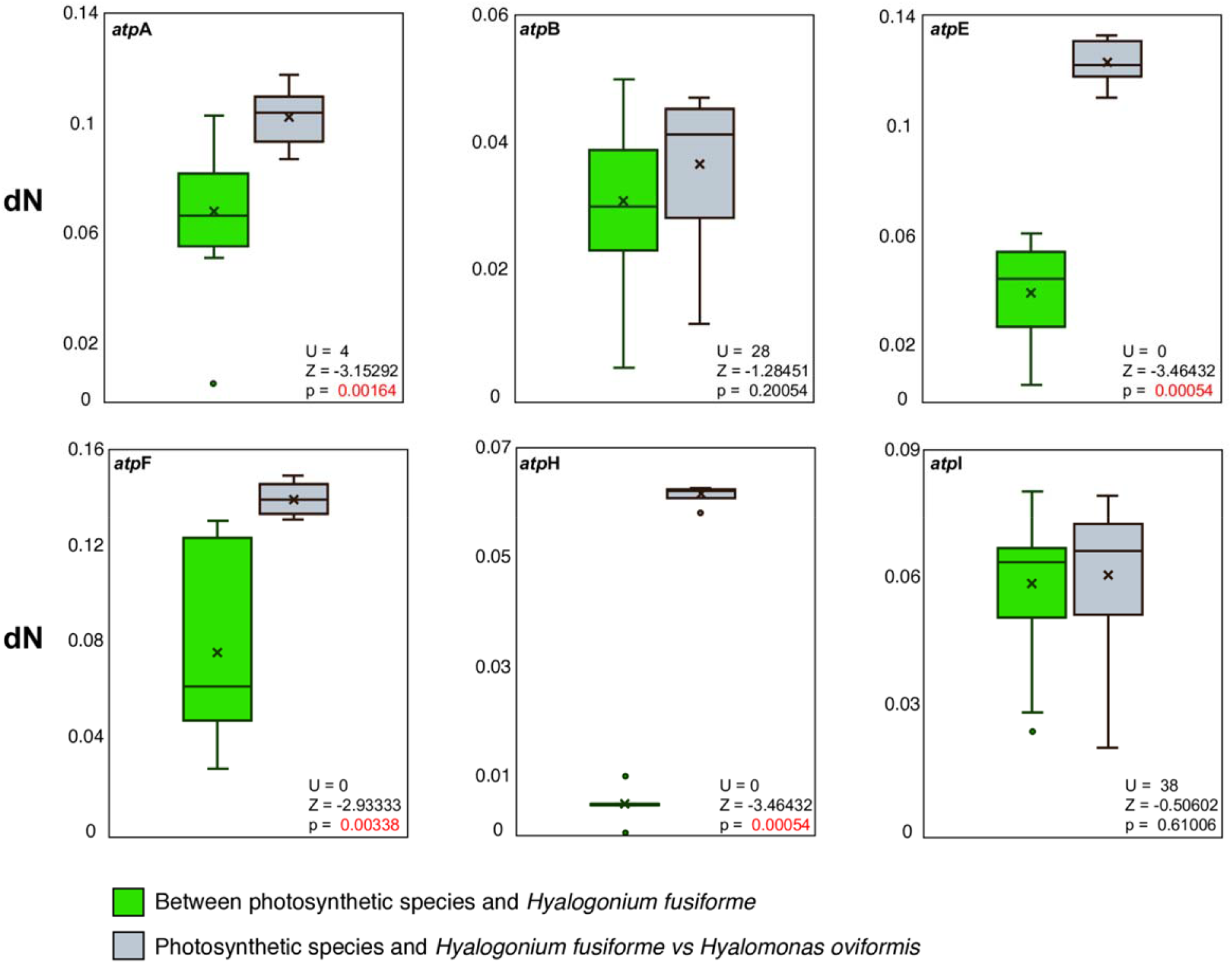
Comparison of nonsynonymous (dN) nucleotide substitution rates estimated in the five plastid genes encoding subunits of the F-ATPase. Box plots in green colour correspond to nucleotide substitution rates estimated between the species *Hm. chlamydogama*, *Chlamydomonas leiostraca*, *Chlamydomonas reinhardtii*, *Chlorogonium capillatum*, *Chlorogonium euchlorum* and *Hyalogonium fusiforme*. Box plots in gray color summarize the nucleotide substitution rates between the former six chlamydomonadalean taxa and *Hyalomonas oviformis*. The *Z* and *U* scores and the *p* values estimated with the Wilcoxon-Mann-Whitney U test are indicated in each case.

The presence of the cF_0_-CF_1_ ATPase in a nonphotosynthetic context is intriguing (Dorrell et al., 2019; Tanifuji et al., 2020). Some have suggested that its hydrolytic activity sustains a proton electrochemical gradient essential for the internalization of proteins into the thylakoid via the twin arginine translocator (TAT) system (Dorrell et al., 2019). Support for this view in *Hm. oviformis* comes from the fact that, in addition to plastid-encoded *atp* genes, we uncovered nuclear genes for *atpD* and *atpG*, which respectively encode the delta and gamma (γ) subunits of the plastid cF_0_-CF_1_ ATPase. The deduced amino acid sequences of *atpD* and *atpG* both contain standard N-terminus plastid transit peptides, and the latter contains the two cysteine residues involved in the thioredoxin-mediated modulation of the enzyme hydrolytic activity (Hisabori et al., 2012) (Figure S7). We also searched the *Hm. oviformis* nuclear contigs for *tat*A, *tat*B, and *tat*C, which encode the TAT protein import system, but only recovered evidence of *tatC*. Similar findings were recently reported in the gene repertoire of *L. pallida*, but in that case *tat*A and *tat*B were identified in transcriptomic data (Pánek et al., 2022).

At circa 463 kb, the ptDNA of *Hg. fusiforme* is not only larger than that of its close photosynthetic relative *Cg. euchlorum* (314 kb; intron-rich genome), but also the largest plastome ever observed in a colorless plant or alga, and larger than the vast majority of sequenced ptDNAs from photosynthetic species. This expanded architecture is primarily the consequence of inflated intergenic regions as well as apparent accumulation of intron-resembling sequences (Tables 1 and S6). Close inspection of the intergenic ptDNA of *Hg. fusiforme* revealed dozens of TrRnEn-like ORFs belonging to different families of homing endonucleases, including members of the LAGLIDADG (2), GIY-YIG (4) and H-N-H (14) families as well as ORFs matching to reverse transcriptases and maturases (see Table S6). These ORFs were not evidently associated with introns and were one of the key reasons why we were unable to bridge the scaffolds during ptDNA assembly. Other chlamydomonadalean algae, including *Dunaliella salina*, also have a preponderance of free-standing intronic ORFs within their intergenic regions (Smith et al., 2010).

**Table 1.** Plastid genome features of diverse chlorophytes.

| Taxa | Genbank Accession | Size (kb) | Inverted repeat length (kb) | Coding DNA (%) | GC content (%) | Mean intergenic spacer (bp) | Protein-coding genes | rRNA genes | tRNA genes | Introns |
| --- | --- | --- | --- | --- | --- | --- | --- | --- | --- | --- |
| **Chlorophyceae** |  |  |  |  |  |  |  |  |  |  |
| *Hyalomonas oviformis*<sup>1</sup> | pending | 132 | NA | 40.6 | 27.3 | 617 | 33 | 5 | 27 | 3 |
| *Hyalomonas chlamydogama*<sup>1</sup> | pending | 198 | 12.1 | 40.9 | 27.6 | 886 | 66 | 10 | 29 | 6 |
| *Hyalogonium fusiforme*<sup>1</sup> | pending | ~442 | NA | 18.4 | 44 | 2454 | 66 | 6 | 21 | 4 |
| *Chlorogonium euchlorum*<sup>1</sup> | pending | ~314 | 27.9 | 27.9 | 36.9 | 1692 | 67 | 6 | 31 | 14 |
| *Chlorogonium capillatum* | KT625085.1-KT620591.1 | ~271 | NA | 34.7 | 38.8 | 1616 | 65 | 3 | 27 | 10 |
| *Polytoma uvella* | KX828177.1 | ~230 | NA | 26.8 | 27.3 | 2967 | 24 | 2 | 27 | 1 |
| *Chlamydomonas leiostraca* | NC_032109 | 167 | 13.1 | 52.4 | 42.5 | 600 | 68 | 6 | 28 | 8 |
| *Volvocales sp. NrCl902* | LC516060.1 | 176 | NA | 39.2 | 40.1 | 1974 | 24 | 5 | 30 | 5 |
| *Chlamydomonas reinhardtii* | NC_005353.1 | 204 | 22.2 | 37.8 | 34.5 | 700 | 68 | 10 | 29 | 6 |
| *Volvox carteri* | GU084820.1 | >420 | 16.6 | <20 | 42.8 | 5103 | 60 | 4 | 27 | 8 |
| *Dunaliella salina* | GQ250045.1 | 269 | 14 | 45.5 | 32.1 | 935 | 66 | 3 | 28 | 43 |
| *Haematococcus pluvialis* | MG677935.1 | 1352 | NA | ~90 | ~50 | NA | ~71-125 | 6 | 31 | NA |
| *Leontynka pallida* | OM479425.1 | 362 | NA | 18 | 36.1 | 4726 | 32 | 3 | 26 | 4 |
| **Trebouxiophyceae** |  |  |  |  |  |  |  |  |  |  |
| *Helicosporidium sp.* | NC_008100.1 | 37 | NA | 95.2 | 26.9 | 34 | 21 | 3 | 25 | 1 |
| *Prototheca wickerhamii* | KJ000176.1 | 56 | NA | 83.17 | 31.2 | 135 | 34 | 3 | 26 | 0 |

The large size of *Hg. fusiforme* ptDNA is not only associated with increased intergenic and intronic regions. In contrast to the ptDNAs of *Hm. oviformis* and other nonphotosynthetic chlamydomonadalean species, the *Hg. fusiforme* plastome shows virtually no gene loss as compared to its photosynthetic close relatives. Indeed, it encodes most subunits of both photosystems (*psa* and *psb*), the *b_6_f* complex, and the proteins for chlorophyll biosynthesis present in *Chlorogonium* species (Table S4). What is more intriguing is that the vast majority of the *Hg. fusiforme* plastid genes typically involved in photosynthesis display no obvious reductions in size or classic signs of pseudogenization (Table S5), with the exception *chl*B (1,068 bp), which is around 130 amino acids shorter than its *Cg. elongatum* counterpart. Still, the *Hg. fusiforme chl*B encodes all the amino acid residues important for the activity of the protochlorophyllide oxidoreductase (Bröcker et al., 2010) (Figure S8). It is notable as well that *psb*E and *psb*Z from *Hg. fusiforme* are split into two pieces by an intergenic spacer (no intron was detected), but their conceptual translations reveal protein products of similar lengths to those of photosynthetic relatives. Evolutionary rate analyses indicate that some *Hg. fusiforme* plastid-encoded proteins involved in light reactions and carbon fixation have higher rates of substitution than their homologs in photosynthetic taxa (Figure S6). But, altogether, there is no clear evidence of rampant plastid pseudogenization in *Hg. fusiforme*, despite it having forfeited photosynthetic capabilities (i.e., no chlorophyll presence; Figure 2A; Pringsheim 1963).

### Plastome evolution following the loss of photosynthesis

The loss of photosynthesis has traditionally been associated with plastid gene loss and genomic streamlining. In parallel, plastome gene repertoires of nonphotosynthetic plants and algae, regardless of their trophic strategy, tend to converge to relatively homogeneous small gene sets involved in transcription and translation (Figure S9). Only in rare cases, the plastome remodeling process after the loss of photosynthesis resulted in the complete loss of the organelle genome (Smith & Lee, 2014). But as more and more ptDNA sequences from colorless species become available, it is increasingly obvious that there is not a single narrative for plastome evolution following the loss of photosynthetic capabilities. This is certainly true for colorless members of the Chlamydomonadales, where plastid genome information exists for at least six nonphotosynthetic lineages, each representing an independent loss of photosynthesis within the order. A key theme emerging from these data is that the ptDNAs of colorless plastids, in particular those of free-living Chlamydomonadales, are not always characterized by small size, reduced gene content and compactness. In fact, diverse combinations of genome size, gene content, and gene density exist in this green algal group. The notable cases include the large gene-rich plastome of *Hg. fusiforme* as well as the large but gene-reduced ptDNAs of *Leontynka pallida* (Pánek et al., 2022) and *Polytoma uvella* (Figueroa-Martinez et al., 2017); nevertheless, all three of these plastid genomes are significantly larger than those of their close photosynthetic relatives. But this group of colorless algae also includes the relatively small and compact plastomes of *Hm. oviformis* and *Volvocales* sp. NrCl902 and the complete loss of ptDNA in *Polytomella* species.

The observed diversity among ptDNAs of colorless chlamydomonadalean algae highlights the fact that various aspects of genome organization – size, compaction, and gene content – evolve independently, even under seemingly similar selective pressures associated with the switch to a strictly heterotrophic lifestyle. This is because they are likely driven by different evolutionary mechanisms and forces. For instance, gene loss is primarily a consequence of (1) reduced functional constraints on photosynthesis-related genes and (2) evolutionary time, whereby the longer the time since the loss of photosynthesis the greater the potential for gene loss. Conversely, changes in genome compaction are largely driven by mutational forces and DNA maintenance machineries acting upon intergenic and intronic regions (Figueroa-Martinez et al., 2015; Wicke et al., 2013). Thus, it is possible for a plastid genome to be large and gene poor, and vice versa.

Our data suggest that *Hg. fusiforme* may have lost photosynthetic capabilities in recent evolutionary time, thus, explaining why it has a near-complete cohort of photosynthetic genes. In the future, it is possible/expected that some of these genes will be lost, but its propensity for genomic expansion may (or may not) persist. It should be noted that chlamydomonadalean ptDNAs, including those of photosynthetic species, are particularly prone to genomic inflation, a topic that has been discussed in detail elsewhere (Smith et al., 2010; Figueroa-Martinez et al., 2017; Gaouda et al., 2018). In fact, the largest plastid genome on record belongs to *Haematococcus pluvialis* (>1 Mb) (Gaouda et al., 2018), a relatively close relative of *Hg. fusiforme*. As it stands, the data presented here add to the astonishing amount of plastome architectural diversity observed across the Chlamydomonadales (in both photosynthetic and nonphotosynthetic species). How much more diversity remains to be uncovered? Our guess is a lot.

## Materials and Methods

### Growth conditions

*Hyalomonas oviformis* (SAG 62-27) and *Hyalogonium fusiforme* (SAG 62-1c) were grown in ErbsMS media (Pringsheim, 1946) under dark conditions and no shaking at 16 °C. Cultures of *Hm. chlamydogama* (SAG 11-48b) were maintained in standard *Volvox* medium (Kirk et al., 1999) under constant shaking (200 rpm) at 18°C and a 16-hour-light/8-hour-dark cycle. *Cg. euchlorum* (SAG 12-2a) cultures were grown in MiEg 1:1 and Eg media (Schlösser, 1994).

### DNA extraction and sequencing

Total DNA of *Hm. chlamydogama* was extracted from a 250-mL culture in exponential growth. The cell pellet was washed three times with saline-EDTA (50 mM Tris-HCl, pH 8, 50 mM NaCl, and 5 mM EDTA) and then incubated in the presence of proteinase K (33 mg/mL) and 0.5% SDS for 1 h at 50°C. DNA was extracted by standard phenol-chloroform procedures. Total genomic DNA from *Hm. oviformis, Hg. fusiforme* and *Cg. euchlorum* was extracted using the DNeasy PlantMini Kit (Qiagen, Hilden, Germany) following manufacturer instructions. Paired-end libraries were sequenced using Illumina technology (150 bp length reads; 450-bp insert length; Genome Quebec, McGill University). We also produced low-coverage PacBio reads (PacBio Sequel, v3 chemistry; IMR Dalhousie University) of *Hg. fusiforme* (413 219 reads; circa 604 Mb) and *Cg. euchlorum* (555 390 reads; circa 680 Mb) from selected DNA fragments of ~ 350 Kbp. The 18S rRNA and ITS rRNA sequences of the used strains are available in GenBank (SAG 62-27: OM985702; SAG 11-48b: OM985703; SAG 62-1c: OM985704; SAG 12-2a: OM985705).

### Genome assembly and sequence annotation

After quality control filtering and trimming, we assembled two high-quality paired-end read sets of each alga alternatively with Ray v 2.2.0 [k-mer lengths of 21, 31, and 77] (Boisvert et al., 2010). PacBio SMRT reads of *Hg. fusiforme* and *Cg. euchlorum* were corrected and assembled with Canu v 2.1 [predicted genome size of 0.4 megabases, using raw PacBio reads for correction and then corrected reads for assembly] (Koren et al., 2017).

We identified contigs containing typical plastid genes using the Automatic Annotation tool of Geneious Prime v2019.1.3 (Kearse et al., 2012) with published complete ptDNAs from diverse Chlamydomonadales as reference. Manual sequence refinement and read coverage evaluations were carried out with Geneious Prime. Genes encoding tRNAs were predicted with the tRNAscan-SE Search Server (Lowe & Chan, 2016).

### Phylogenetics, relative divergence time and nucleotide substitution rates analyses

Sequence multiple alignments for all analyses were prepared alternatively with the MAFFT (Katoh & Standley, 2013) tools implemented in Geneious Prime. Resulting raw sequence matrices were visually inspected and manually edited to discard poorly aligned regions. To test the independent origin of the colourless lineages *Hm. oviformis* and *Hg. fusiforme* analyzed here, we estimated maximum likelihood (ML) trees from multiple alignments of nuclear 18S rRNA (1,521 bp long; 121 taxa) and plastid 16S rRNA (1,257 bp long and 35 taxa) sequences of diverse chlorophyte green algae. Both ML reconstruction were estimated independently with IQ-TREE v1.6.11 (Nguyen et al., 2015) considering the best fitting nucleotide substitution model (TIM3+R3 and TVMe+I+G4, selected respectively by the ModelFinder version implemented in IQ-TREE v1.6.11) and 10,000 ultra-fast bootstrap replicates in each case. Then, to estimate the relative divergence times (rDT) of the Chlamydomonadalean colorless lineages, we prepared additional alignments of 16S rRNA+18S rRNA (2,763 nucleotides long; 28 taxa) sequences and of conceptual translations of the protein-coding genes *tufa, rps*4, *rps*7, *rpl*2 and *rpl*5 (1,163 amino acid residues long; 28 taxa) considering a reduced sample of chlorophyte algae (see supplementary Table S7). The five protein-coding sequences were selected based of their presence in ptDNAs of the colorless algae analyzed here, the length of the translated protein (>150 aa) and reliability (i.e., limited ambiguously aligned or gap-rich regions) of the multiple alignments. Maximum likelihood trees of the 16S rRNA, 18S rRNA, the concatenated 16S rRNA and 18S rRNA sets and concatenated protein alignments were independently estimated with IQ-TREE considering the best fitting model for each sequence matrix (-m TEST option). Then, we used those maximum likelihood phylogenetic trees as references to estimate rDT with RelTime approximation implemented in MEGA X (Mello, 2018). We considered the *Proterocladus* fossils (780 −1000 MY) as the minimum divergence time between Ulvophyceae-Briopsidales and Chlorophyceae as calibration reference point (Tang et al., 2020).

Finally, we also prepared multiple alignments at nucleotide sequence level of selected protein-coding regions from diverse photosynthetic and free-living non-photosynthetic chlamydomonadalean algae to estimate rates of synonymous (*dS*) and nonsynonymous (*dN*) substitutions with the CODEML option of PAMLX (Xu & Yang, 2013).

## Accession numbers

Data deposition: sequences have been deposited in GenBank under accession numbers XXXXX

## Acknowledgments

This work was supported by Discovery Grant to A.R.P. from the Natural Sciences and Engineering Research Council (NSERC) of Canada. A.R.P. was also supported by the Canada Foundation for Innovation (project 28276) and the New Brunswick Innovation Foundation (project RIF2012-006).

## Supplemental data

### Supplemental tables

**Supplemental Table S1.** RelTime estimated divergence times of non-photosynthetic chlamydomonadalean algae

**Supplemental Table S2.** General information of the Illumina and PacBio assemblies

**Supplemental Table S3**. Average read coverage estimated per genomic compartment

**Supplemental Table S4**. Comparative gene content of plastid genomes from diverse chlamydomonadalean algae

**Supplemental Table S5**. Size of predicted plastid protein-coding regions in *Hyalogonium*

**Supplemental Table S6.** Functional classification of the plastid *Hyalogonium fusiforme* TrRnEn-like ORFs

**Supplemental Table S7**. Taxa considered in the RelTime analyses

### Supplemental Figures

**Supplemental Figure S1**. Divergence time estimates of nonphotosynthetic lineages among the Order Chlamydomonadales based on the *RelTime* approximation (Tamura et al., 2012) applied to **A**) an alignment of 5 plastid encoded proteins (*tuf*A, *rps*4, *rps*7, *rpl*2 and *rpl*5) and **B**) a concatenated alignment of the 16SrRNA (plastid) and 18S rRNA (nuclear) sequences. For illustration purposes, the presented trees correspond to subsections of the highest-likelihood trees estimated with IQ-TREE in each case (LG+R4 for the protein set and GTR+R3 for the concatenated rRNA data). The complete rooted trees including the outgroup (*Chlorella* clade) and indicating the fossil calibration point (*Proteolcadus* fossils; 780 −1000 MY) are available in the supplementary information (Figs. S4 and S5). Black numbers near nodes indicate values of bootstrap support (10,000 ultra-fast replicates) and red numbers are the *RelTime* estimated divergence times. Yellow rectangles delimit 95% credibility intervals on node ages. The branch lengths are proportional to the absolute ages of the nodes.

**Supplemental Figure S2**. Linear maps of the four assembled scaffolds of the *Chlorogonium euchlorum* ptDNA. The graphic representations were created with OGDraw (Greiner et al., 2019).

**Supplemental Figure S3.** Alignment of the complete plastid genomes of *Hyalomonas oviformis* and *Hm. chlamydogama*. Colored diagonal lines indicate conserved gene blocks. Rectangles in red highlight protein-coding genes associated to the photosynthetic function. Grey rectangles indicate genes encoding rRNAs. The genome alignment was elaborated with the MAUVE v 2.4.0 tool (progressive algorithm default options) implemented in Geneious Prime.

**Supplemental Figure S4.** Divergence time estimates of nonphotosynthetic lineages among the Order Chlamydomonadales based on a RelTime approximation (Tamura et al., 2012) applied to an alignment of 5 plastid encoded proteins (*tuf*A, *rps*4, *rps*7, *rpl*2 and *rpl*5). The tree topology corresponds to a the highest-likelihood tree (Log Likelihood = −16540.505) estimated with IQ-TREE considering the best fitting amino acid substitution model (LG+R4) identified by the ModelFinder version implemented in IQ-TREE v1.6.11. Nonphotosynthetic taxa are highlighted with orange circles to the right. The outgroup (*Chlorella* clade) and the fossil calibration point (*Proteolcadus* fossils; 780 −1000 MY; red dot) are indicated. Black numbers near nodes indicate values of bootstrap support (10,000 ultra-fast replicates) and red numbers are the *RelTime*-estimated divergence times. Grey rectangles delimit 95% credibility intervals on node ages. The branch lengths proportional to the absolute ages of the nodes.

**Supplemental Figure S5.** Divergence time estimates of nonphotosynthetic lineages among the Order Chlamydomonadales based on a RelTime approximation (Tamura et al., 2012) applied to a concatenated alignment of the 16SrRNA and 18S rRNA sequences. The tree topology corresponds to a the highest-likelihood tree (Log Likelihood = −14261.073) estimated with IQ-TREE considering the best fitting model (GTR+R3) identified by identified by the ModelFinder version implemented in IQ-TREE v1.6.11. Nonphotosynthetic taxa are highlighted with orange circles to the right. The outgroup (*Chlorella* clade) and the fossil calibration point (*Proteolcadus* fossils; 780 −1000 MY; red dot) are indicated. Black numbers near nodes indicate values of bootstrap support (10,000 ultra-fast replicates), red numbers are the *RelTime*-estimated divergence times. Grey rectangles delimit 95% credibility intervals on node ages. The branch lengths proportional to the absolute ages of the nodes.

**Supplemental Figure S6**. Comparison of nonsynonymous (dN) and synonymous (dS) nucleotide substitution rates estimated in photosynthesis-related plastid genes of *Hyalogonium fusiforme*. Box plots in green color correspond to nucleotide substitution rates estimated between the photosynthetic species *Hyalomonas chlamydogama*, *Chlamydomonas leiostraca*, *Chlamydomonas reinhardtii*, *Chlorogonium capillatum* and *Chlorogonium euchlorum*. Box plots colored in gray summarize the nucleotide substitution rates between the five mentioned photosynthetic taxa and *Hyalogonium fusiforme*.

**Supplemental Figure S7.** Multiple protein sequence alignments of the **A**) γ and **B**) δ subunits of the cF_0_-CF_1_ ATPase from diverse viridiplants. Amino acid residues highlighted in yellow colour represent the transit peptides for import into the plastid predicted by TargetP 2.0 [http://www.cbs.dtu.dk/services/TargetP/] (Armenteros et al., 2019). In the case of the γ subunit multiple alignment (panel **A**), the columns emphasized in black background indicate the two cysteine residues typically involved in the thiol-modulation of the cF_0_-CF_1_ ATPase activity.

**Supplemental Figure S8.** Multiple protein sequence alignments of the ChlB from *Chlorogonium* species and *Hyalogonium fusiforme*. Residues highlighted in magenta and cyan represent amino acids involved in the binding and stabilization of the [4Fe-4S] cluster of the protochlorophyllide oxidoreductase. The residues with green background indicate amino acids participating in the interaction of ChlB (binding) with ChlL (Bröcker et al., 2010).

**Supplemental Figure S9.** Comparative content of protein-coding genes in plastid genomes of diverse non photosynthetic green algae and *Euglena (Astasia) longa*, which contains a secondary plastid of green algal origin. Gene names with asterisks refer to *gene series* encoding subunit of the CF-ATPase (*atp* series), the *b_6_f* complex (*pet* series), photosystem I (*psa* series) and photosystem II (*psb* series). Taxa codes: As, *Euglena longa*; Pw, *Prototheca wickerhamii*; He, *Helicosporidium* sp.; Pu, *Polytoma uvella*; Vs, *Volvocales* NrCl902; Ho, *Hyalomonas oviformis*; Hf, *Hyalogonium fusiforme*.

## Parsed Citations

Armenteros JJA, Salvatore, M, Emanuelsson, O, Winther, O, Heijne, G von, Elofsson A, Nielsen H (2019). Detecting sequence signals in targeting peptides using deep learning. Life Science Alliance, 2: e201900429. https://doi.org/10.26508/LSA.201900429

Boisvert, S, Laviolette, F, Corbeil J (2010) Ray: Simultaneous assembly of reads from a mix of high-throughput sequencing technologies. J Comput Biol 17: 1519–1533. https://doi.org/10.1089/cmb.2009.0238

Bold HC (1949). The morphology of Chlamydomonas chlamydogama, sp. nov. Bull Torrey Bot Cl., 76: 101–108. https://doi.org/10.2307/2482218

Bröcker MJ, Schomburg, S, Heinz, DW, Jahn, D, Schubert, WD, Moser J (2010). Crystal structure of the nitrogenase-like dark operative protochlorophyllide pxidoreductase catalytic complex (ChlN/ChlB)2. J Biol Chem 285: 27336–27345. https://doi.org/10.1074/jbc.M110.126698

Dorrell, R.G., Azuma T, Nomura M, de Kerdrel GA, Paoli L, Yang S, Bowler C, Ishii K, Miyashita H, Gile GH, Kamikawa R (2019) Principles of plastid reductive evolution illuminated by nonphotosynthetic chrysophytes. Proc Natl Acad Sc USA 116: 6914–6923. https://doi.org/10.1073/pnas.1819976116

Figueroa-Martinez F, Nedelcu AM, Smith DR, Reyes-Prieto A (2015). When the lights go out: the evolutionary fate of free-living colorless green algae. New Phytologist 206: 972–982. https://doi.org/10.1111/nph.13279

Figueroa-Martinez F, Nedelcu AM, Smith DR, Reyes-Prieto A (2017) The plastid genome of Polytoma uvella is the largest known among colorless algae and plants and reflects contrasting evolutionary paths to nonphotosynthetic lifestyles. Plant Physiol 173: 932–943. https://doi.org/10.1104/pp.16.01628

Gaouda H, Hamaji T, Yamamoto, K, Kawai-Toyooka H, Suzuki M, Noguchi H, Minakuchi Y, Toyoda A, Fujiyama A, Nozaki H, Smith D. R. (2018). Exploring the limits and causes of plastid genome expansion in Volvocine green algae. Genome Biol Evol 10: 2248–2254. https://doi.org/10.1093/GBE/EVY175

Greiner S, Lehwark P, Bock R (2019) OrganellarGenomeDRAW (OGDRAW) version 1.3.1: expanded toolkit for the graphical visualization of organellar genomes. Nucleic Acids Res 47: W59–W64. https://doi.org/10.1093/nar/gkz238

Hisabori T, Sunamura EI, Kim, Y, Konno H (2012). The chloroplast ATP synthase features the characteristic redox regulation machinery. Antioxid Redox Signal 19: 1846–1854. https://doi.org/10.1089/ars.2012.5044

Katoh K, Standley DM (2013) MAFFT Multiple Sequence Alignment Software Version 7: Improvements in performance and psability. Mol Biol Evol 30: 772–780. https://doi.org/10.1093/MOLBEV/MST010

Kearse M, Moir R, Wilson A, Stones-Havas S, Cheung M, Sturrock S, Buxton S, Cooper A, Markowitz S, Duran C, Thierer T, Ashton B, Meintjes P, Drummond A (2012). Geneious Basic: An integrated and extendable desktop software platform for the organization and analysis of sequence data. Bioinformatics 28: 1647–1649. https://doi.org/10.1093/bioinformatics/bts199

Kirk MM, Stark K, Miller SM, Muller W, Taillon BE, Gruber H, Schmitt R, Kirk DL (1999) regA, a Volvox gene that plays a central role in germ-soma differentiation, encodes a novel regulatory protein. Development 126: 639–647. https://dev.biologists.org/content/126/4/639

Kayama M, Chen JF, Nakada T, Nishimura Y, Shikanai T, Azuma T, Miyashita H, Takaichi S, Kashiyama Y, Kamikawa R (2020) A non-photosynthetic green alga illuminates the reductive evolution of plastid electron transport systems. BMC Biology 18: 126. https://doi.org/10.1186/s12915-020-00853-w

Koren S, Walenz BP, Berlin K, Miller JR, Bergman NH, Phillippy AM (2017) Canu: scalable and accurate long-read assembly via adaptive k-mer weighting and repeat separation. Genome Res 27: 722–736 https://doi.org/10.1101/gr.215087.116

Lowe TM, Chan PP (2016) tRNAscan-SE On-line: integrating search and context for analysis of transfer RNA genes. Nucleic Acids Res 44: W54–57. https://doi.org/10.1093/nar/gkw413

Mello B (2018). Estimating TimeTrees with MEGA and the TimeTree Resource. Mol Biol Evol 35: 2334–2342. https://doi.org/10.1093/molbev/msy133

Nedelcu AM (2001) Complex patterns of plastid 16S rRNA gene evolution in nonphotosynthetic green algae. J Mol Evol 53: 670–679. https://doi.org/10.1007/s002390010254

Nguyen LT, Schmidt HA, von Haeseler A, Minh BQ (2015) IQ-TREE: A fast and effective stochastic algorithm for estimating maximum-likelihood phylogenies. Mol Biology Evol 32: 268–274. https://doi.org/10.1093/molbev/msu300

Pánek T, Barcytė D, Treitli SC, Záhonová K, Sokol M, Ševčíková T, Zadrobílková E, Jaške K, Yubuki N, Čepička I, Eliáš M (2022). A new lineage of non-photosynthetic green algae with extreme organellar genomes. BMC Biology 20: 66 https://doi.org/10.1186/s12915-022-01263-w

Pringsheim EG (1946). Pure Cultures of Algae: their preparation and maintenance. Cambridge University Press, UK.

Pringsheim EG (1963). Farblose Algen. Ein Beitrag zur Evolutionsforschung. Gustav Fischer Verlag, Stuttgart, Germany.

Schlösser UG (1994) SAG-Sammlung von Algenkulturen at the University of Göttingen. Catalogue of strains. Bot Acta 107: 113186. https://doi.org/10.1111/j.1438-8677.1994.tb00784.x

Smith DR, Lee RW (2014) A Plastid without a Genome: Evidence from the Nonphotosynthetic Green Algal Genus Polytomella. Plant Physiol 164: 1812–1819. https://doi.org/10.1104/pp.113.233718

Smith DR, Lee RW, Cushman JC, Magnuson JK, Tran D, Polle JEW (2010). The Dunaliella salina organelle genomes: large sequences, inflated with intronic and intergenic DNA. BMC Plant Biol 10: 83. https://doi.org/10.1186/1471-2229-10-83

Tamura K, Battistuzzi, FU, Billing-Ross P, Murillo O, Filipski A, Kumar S (2012) Estimating divergence times in large molecular phylogenies. Proc Natl Acad Sc USA 109: 19333–19338. https://doi.org/10.1073/pnas.1213199109

Tang Q, Pang K, Yuan, X, Xiao S (2020) A one-billion-year-old multicellular chlorophyte. Nat. Ecol Evol 4: 543–549. https://doi.org/10.1038/s41559-020-1122-9

Tanifuji G, Kamikawa R, Moore CE, Mills T, Onodera NT, Kashiyama, Archibald JM, Inagaki Y, Hashimoto T (2020) Comparative plastid genomics of Cryptomonas species reveals fine-scale genomic responses to loss of photosynthesis. Genome Biol Evol 12: 3926–3937. https://doi.org/10.1093/gbe/evaa001

Vernon D, Gutell RR, Cannone JJ, Rumpf RW, Birky J (2001) Accelerated evolution of functional plastid rRNA and elongation factor genes due to reduced protein synthetic load after the loss of photosynthesis in the chlorophyte alga Polytoma. Mol Biol Evol 18: 1810–1822. https://doi.org/10.1093/oxfordjournals.molbev.a003968

Wicke S, Müller KF, de Pamphilis CW, Quandt D, Wickett NJ, Zhang Y, Renner SS, Schneeweiss GM (2013) Mechanisms of functional and physical penome peduction in photosynthetic and nonphotosynthetic parasitic plants of the broomrape family. Plant Cell 25: 3711–3725. https://doi.org/10.1105/tpc.113.113373

Xu B, Yang Z (2013) PamlX: A graphical user interface for PAML. Mol Biol Evol 30: 2723–2724. https://doi.org/10.1093/molbev/mst179

