## Supplemental Figure S1 for "The plastomes of *Hyalomonas oviformis* and *Hyalogonium fusiforme* evolved dissimilar architecture after the loss of photosynthesis"

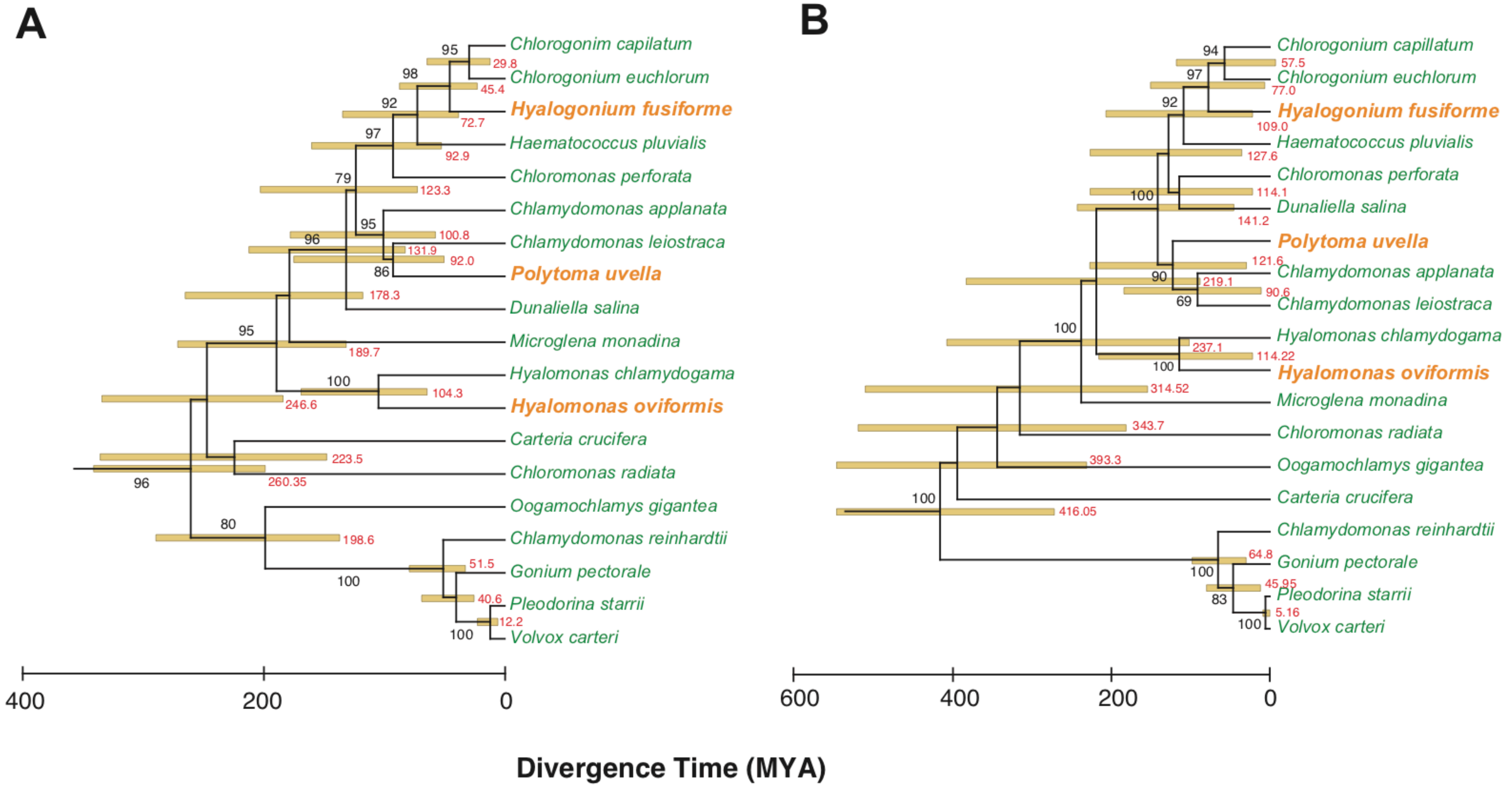


**Supplemental Figure S1**. Divergence time estimates of nonphotosynthetic lineages among the Order Chlamydomonadales based on the *RelTime* approximation (Tamura et al., 2012) applied to **A**) an alignment of 5 plastid encoded proteins (*tuf*A, *rps*4, *rps*7, *rpl*2 and *rpl*5) and **B**) a concatenated alignment of the 16SrRNA (plastid) and 18S rRNA (nuclear) sequences. For illustration purposes, the presented trees correspond to subsections of the highest-likelihood trees estimated with IQ-TREE in each case (LG+R4 for the protein set and GTR+R3 for the concatenated rRNA data). The complete rooted trees including the outgroup (*Chlorella* clade) and indicating the fossil calibration point (*Proteolcadus* fossils; 780 -1000 MY) are available in the supplementary information (Figs. S4 and S5).
