## Supplemental Figure S2 for "The plastomes of *Hyalomonas oviformis* and *Hyalogonium fusiforme* evolved dissimilar architecture after the loss of photosynthesis"

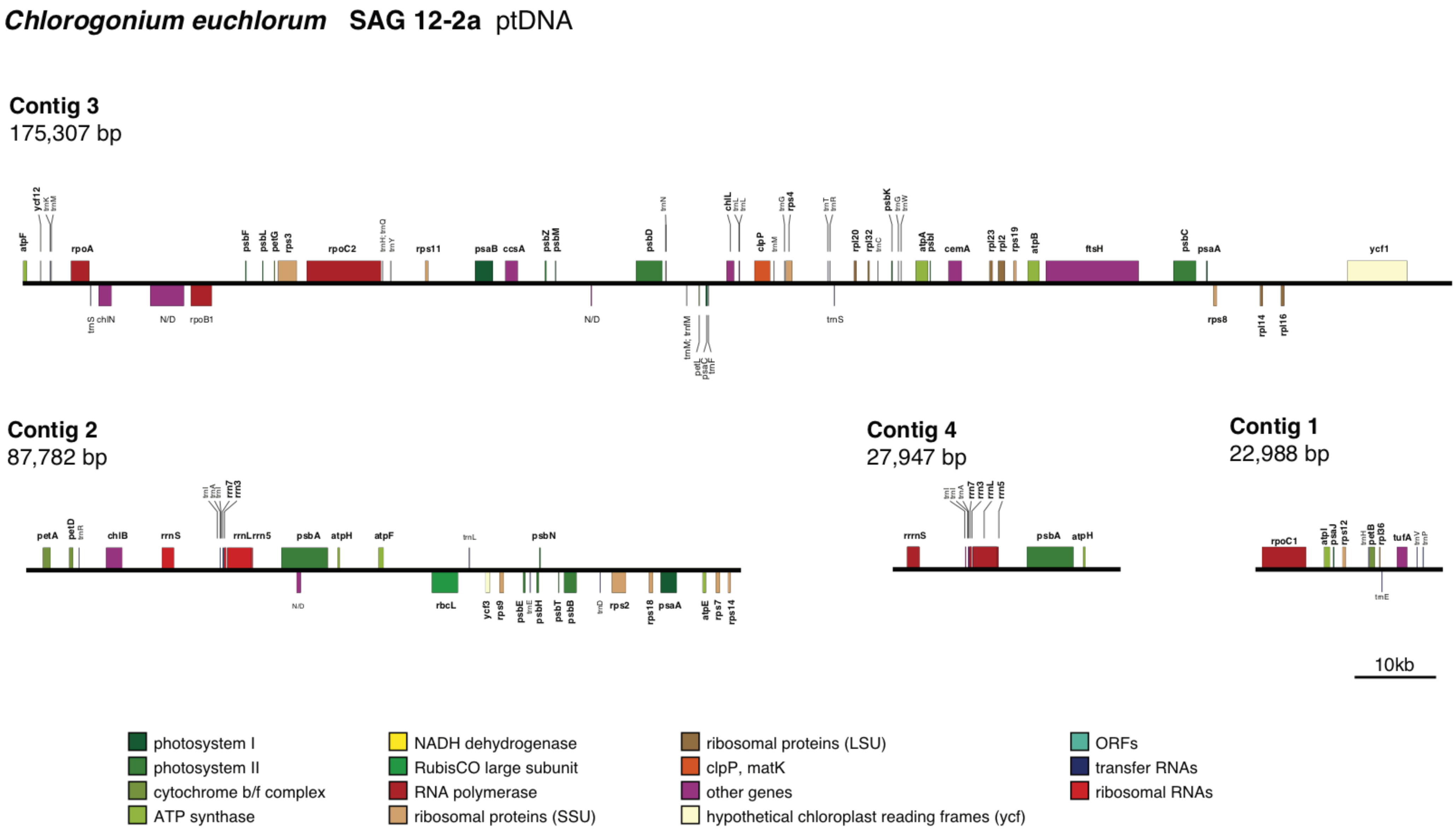


**Supplemental Figure S2**. Linear maps of the four assembled scaffolds of the *Chlorogonium euchlorum* ptDNA. The graphic representations were created with OGDraw (Greiner et al., 2019).
