## Supplemental Figure S3 for "The plastomes of *Hyalomonas oviformis* and *Hyalogonium fusiforme* evolved dissimilar architecture after the loss of photosynthesis"

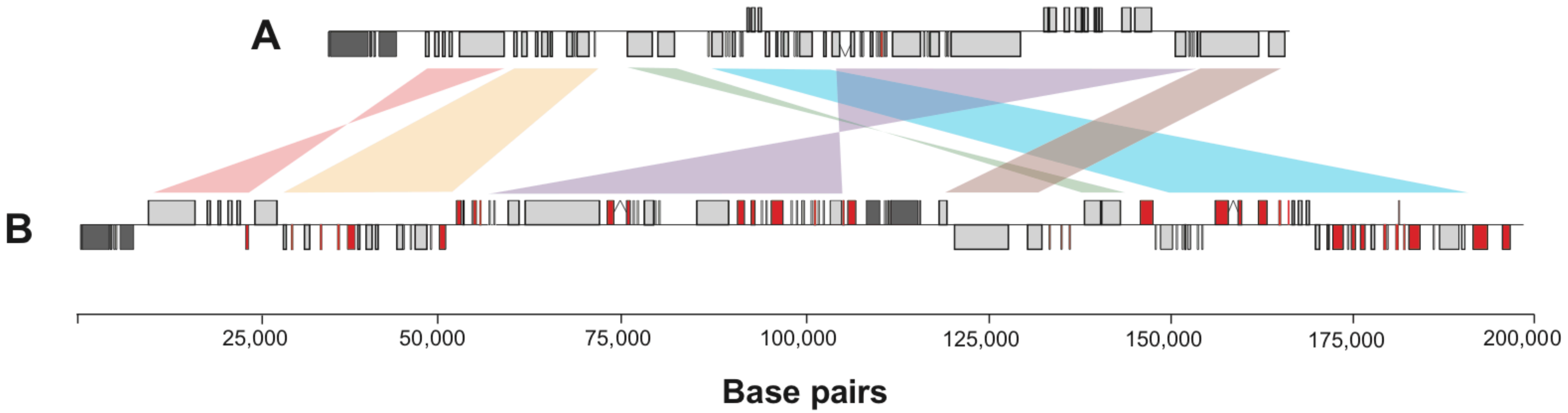


**Supplemental Figure S3**. Alignment of the complete plastid genomes of *Hyalomonas oviformis* and *Hm. chlamydogama*. Colored diagonal lines indicate conserved gene blocks. Rectangles in red highlight protein-coding genes associated to the photosynthetic function. Grey rectangles indicate genes encoding rRNAs. The genome alignment was elaborated with the MAUVE v 2.4.0 tool (progressive algorithm default options) implemented in Geneious Prime.
