## Supplemental Figure S4 for "The plastomes of *Hyalomonas oviformis* and *Hyalogonium fusiforme* evolved dissimilar architecture after the loss of photosynthesis"

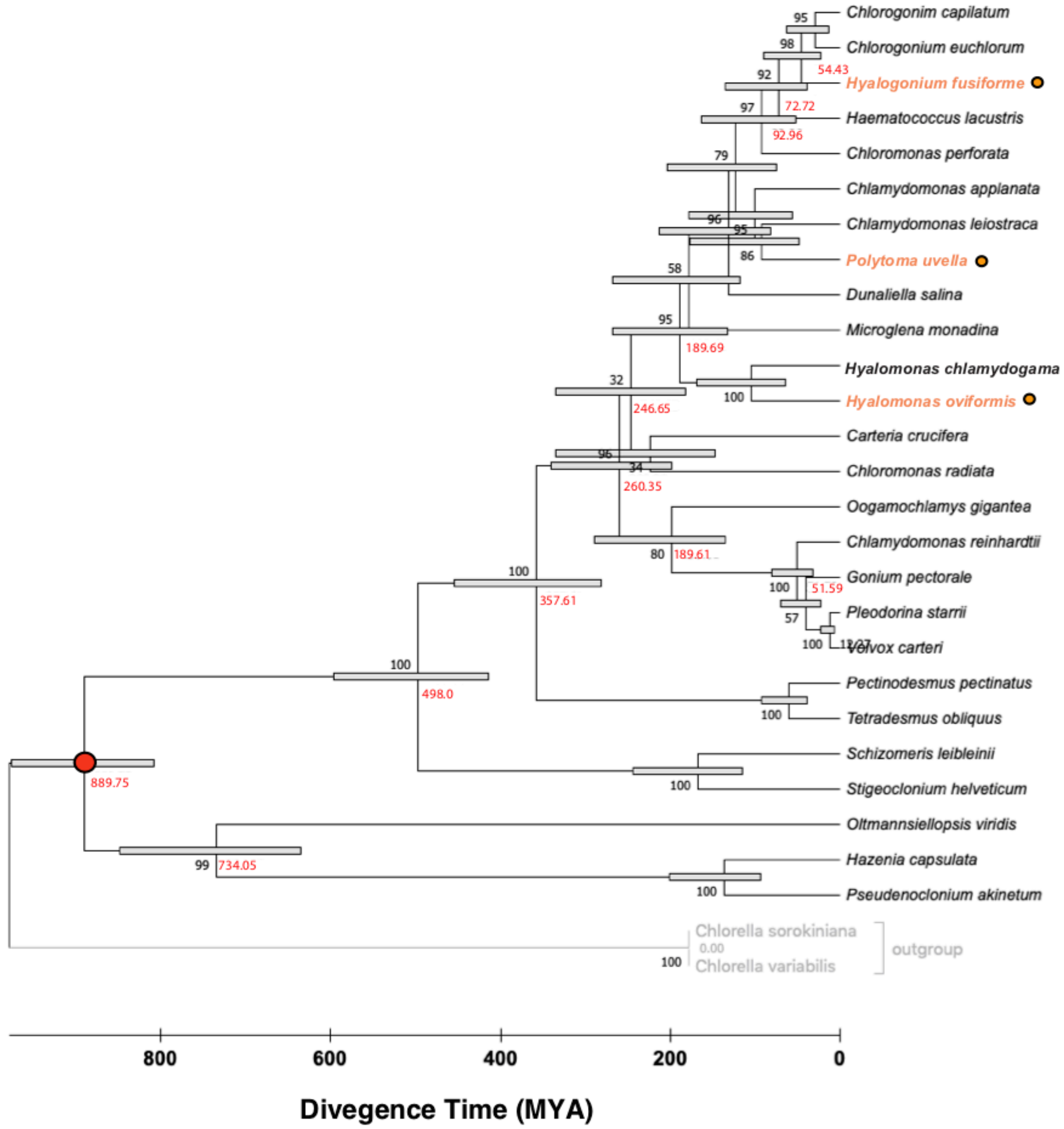


**Supplemental Figure S4**. Divergence time estimates of nonphotosynthetic lineages among the Order Chlamydomonadales based on a *RelTime* approximation (Tamura et al., 2012) applied to an alignment of 5 plastid encoded proteins (*tuf*A, *rps*4, *rps*7, *rpl*2 and *rpl*5). The tree topology corresponds to a the highest-likelihood tree (Log Likelihood = -16540.505) estimated with IQ-TREE considering the best fitting amino acid substitution model (LG+R4) identified by the ModelFinder version implemented in IQ-TREE v1.6.11. Nonphotosynthetic taxa are highlighted with orange circles to the right. The outgroup (*Chlorell*a clade) and the fossil calibration point (*Proteolcadus* fossils; 780 -1000 MY; red dot) are indicated. Black numbers near nodes indicate values of bootstrap support (10,000 ultra-fast replicates) and red numbers are the RelTime-estimated divergence times. Grey rectangles delimit 95% credibility intervals on node ages. The branch lengths proportional to the absolute ages of the nodes.
