## Supplemental Figure S5 for "The plastomes of *Hyalomonas oviformis* and *Hyalogonium fusiforme* evolved dissimilar architecture after the loss of photosynthesis"

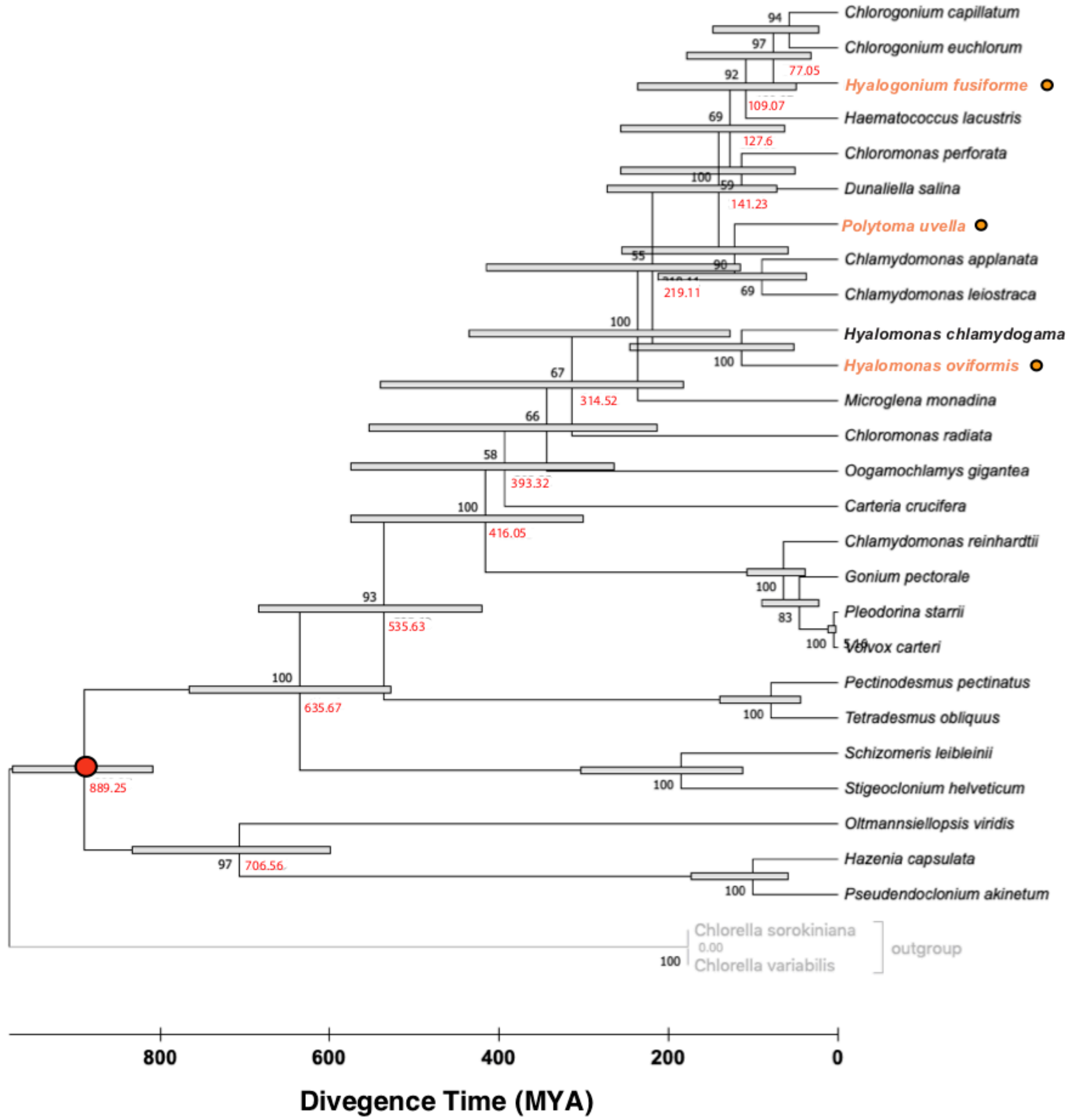


**Supplemental Figure S5**. Divergence time estimates of nonphotosynthetic lineages among the Order Chlamydomonadales based on a *RelTime* approximation (Tamura et al., 2012) applied to a concatenated alignment of the 16SrRNA and 18S rRNA sequences. The tree topology corresponds to a the highest-likelihood tree (Log Likelihood = -14261.073) estimated with IQ-TREE considering the best fitting model (GTR+R3) identified by identified by the ModelFinder version implemented in IQ-TREE v1.6.11. Nonphotosynthetic taxa are highlighted with orange circles to the right. The outgroup (*Chlorella* clade) and the fossil calibration point (Proteolcadus fossils; 780 -1000 MY; red dot) are indicated. Black numbers near nodes indicate values of bootstrap support (10,000 ultra-fast replicates), red numbers are the RelTime-estimated divergence times. Grey rectangles delimit 95% credibility intervals on node ages. The branch lengths proportional to the absolute ages of the nodes.
