## Supplemental Figure S6 for "The plastomes of *Hyalomonas oviformis* and *Hyalogonium fusiforme* evolved dissimilar architecture after the loss of photosynthesis"

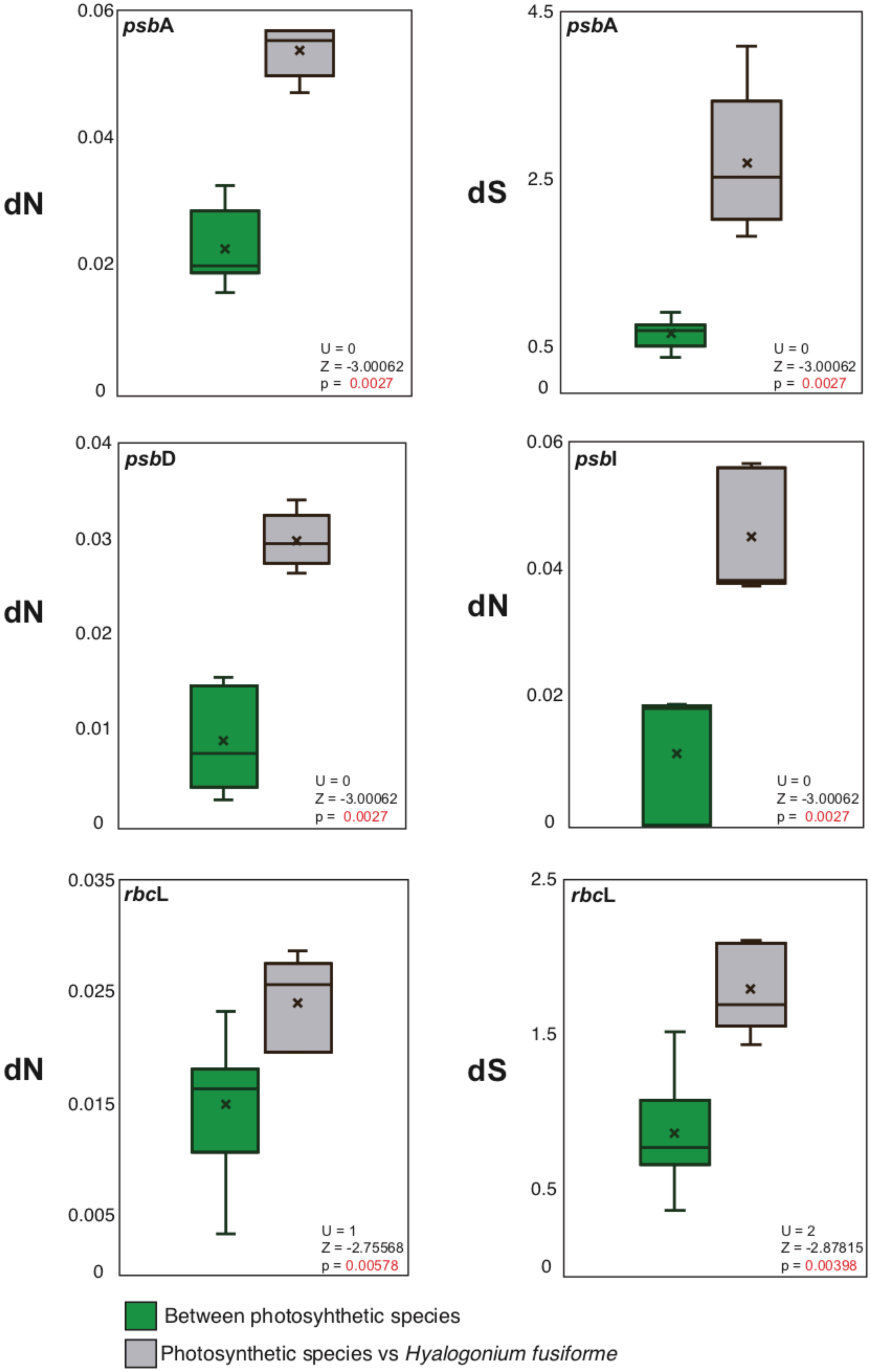


**Supplemental Figure S6**. Comparison of nonsynonymous (dN) and synonymous (dS) nucleotide substitution rates estimated in photosynthesis-related plastid genes of *Hyalogonium fusiforme*. Box plots in green color correspond to nucleotide substitution rates estimated between the photosynthetic species *Hyalomonas chlamydogama, Chlamydomonas leiostraca, Chlamydomonas reinhardtii, Chlorogonium capillatum* and *Chlorogonium euchlorum*. Box plots colored in gray summarize the nucleotide substitution rates between the five mentioned photosynthetic taxa and Hyalogonium fusiforme. The Z and U scores and the p values estimated with the Wilcoxon-Mann-Whitney U test are indicated.
