## Supplemental Figure S7 for "The plastomes of *Hyalomonas oviformis* and *Hyalogonium fusiforme* evolved dissimilar architecture after the loss of photosynthesis"

## A

##### *Hyalomonas oviformis* γ subunit of $cF_OF_1$ ATPase

**TargetP 2.0 prediction**: Thylakoid luminal transfer peptide
CS pos: 35-36. VTA-GL. Pr: 0.7331
Chloroplast transfer peptide likelihood = 0.6874

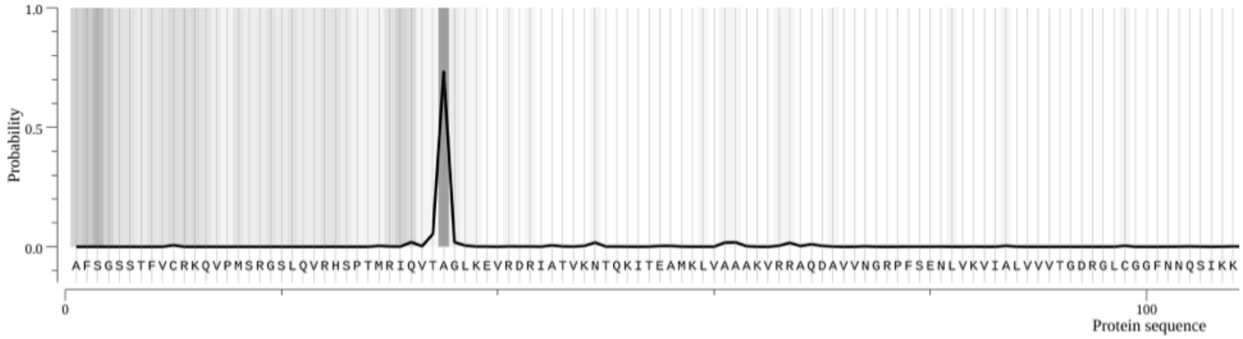


##### Multipe alignment γ subunit of $cF_OF_1$ ATPase

```
A. thaliana 1    MTGSISTSWLLSSPSNSNSASSSESYSFIAT---LKPVRYYPF-----------QSLTPN
A. thaliana 2    M----ACSNLTTMWVSSKPSLSADSSSLSFRSVLKCPT---------------NTSSPPS
S. oleracea      M----ACS-------------LSFSSSVSTFHLPTTTQ---------------STQAPPN
C. braunii       MAAAMATSSAVTAAAAAAGMCSSGSATLSRKKALAAPSTLPSFEGLSASNRLPSRLAPPM
H. lacustris     -----------------MASLMSSKSAFVGR----------------------NASFAPS
C. reinhardtii   M----------------AAMLASKQGAFMGR----------------------SSFAPA
V. carteri       M---------------AATMLASKQGAFMGR----------------------STFAPA
T. sociales      M-------------------------PFGAL----------------------TFDSMS
D. salina        S---------------ALTQFSSKQAAFVGK----------------------AQTCPA
C. eustigma      -----------------MAAISAKQTSFMAS----------------------TSFASS
H. oviformis     -------------------------AFSGS----------------------STFV-C

A. thaliana 1    RI-----SS-RSPLPS----------IQIRAGIRELRERIDSVKNTQKITEAMRLVAAAR
A. thaliana 2    RA-----SS-V---SP----------LQ--ASLRELRDRIDSVKNTQKITEAMKLVAAAK
S. oleracea      NA-----TT-LPTTNP----------IQC-ANLRELRDRIGSVKNTQKITEAMKLVAAAK
C. braunii       SADAELESM-RPETGR----------VTV-CGLKDLRDRIDSVKNTQKITDAMKLVAAAK
H. lacustris     RQ-----QAPVAKRAT----------LQVTAGIKEVRDRIASVKNTMKITEAMKLVAAAK
C. reinhardtii   P------KG-VASRGS----------LQVVAGLKEVRDRIASVKNTQKITDAMKLVAAAK
V. carteri       QQ-----QG-VARRGS----------LQVTAGLKEVRDRIASVKNTNKITDAMKLVAAAK
T. sociales      RC-----QHPSLPACLL---------GDVTAGLKEVRDRIASVKNTMKITDAMKLVAAAK
D. salina        PM-----VA-KPQRGS----------LQVTAGMKEVRDRIGSVKNTQKITEAMKLVAAAK
C. eustigma      KK-----QA-VVQRGA----------LQVSAGMKEIRDRISSVKNTSKITEAMKLVAAAK
H. oviformis     RK-----QV-PMSRGSLQVRHSPTMRIQVTAGLKEVRDRIATVKNTQKITEAMKLVAAAK
                                               ..::::*:** :**** ***:**:*****:
```

### Figure S7

```
A. thaliana 1     VRRAQDAVIKGRPFTETLVEILYSINQSAQLEDIDFPLSIVRPVKRVALVVVTGDKGLCG
A. thaliana 2     VRRAQEAVVNGRPFSETLVEVLYNINEQLQTDDVDVPLTKVRPVKKVALVVVTGDRGLCG
S. oleracea       VRRAQEAVVNGRPFSETLVEVLYNMNEQLQTEDVDVPLTKIRTVKKVALMVVTGDRGLCG
C. braunii        VRRAQEAVVGARPFSENLVKVLYGVNQQIQTEDIDIPLTTVRPVKKIALVVVTGDRGLCG
H. lacustris      VRRAQEAVINSRPFTENLIKVLFGVNQRLRTEDVDSPLCSIRPVKSVLLVVLTGDRGLCG
C. reinhardtii    VRRAQEAVVNGRPFSENLVKVLYGVNQRVRQEDVDSPLCAVRPVKSVLLVVLTGDRGLCG
V. carteri        VRRAQEAVVNGRPFSENLVKVLYGVNQRIRQEDVDSPLCAVRPVKSVMLVVLSGDRGLCG
T. sociales       VRRAQEAVVNGRPFSENLVKVLYGVNQRVRQEDVDSPLCSVRPVKSVMVVVLTGDRGLCG
D. salina         VRRAQDAVINSRPFTENLVKVLYGVNQRVRVEDVDSPLCEIRPVQTVQLVCMTGDRGLCG
C. eustigma       VRRAQEAVINSRPFSENLVKVLYGVNQRLRVEDVDSPLCNIRPVKTVMLVVITGDRGLCG
H. oviformis      VRRAQDAVVNGRPFSENLVKV-------------------------IALVVVTGDRGLCG
                  *****:**: .***:*.*:::                         : :: ::**:****

A. thaliana 1     GFNNAVTKKATLRVQELKQRGIDCVVISVGKKGNAYFSRRDEFDVDKCIEGGGVFPTTKE
A. thaliana 2     GFNNFIIKKAEARIKELKGLGLEYTVISVGKKGNSYFLRRPYIPVDKYLEAGTL-PTAKE
S. oleracea       GFNNMLLKKAESRIAELKKLGVDYTIISIGKKGNTYFIRRPEIPVDRYFDGTNL-PTAKE
C. braunii        GFNNFVLKKAEERRGQIEQMGLECTVVSVGKKGNAFYKRRPQIPLDRTLECGGA-PTTKE
H. lacustris      GYNNFILKKVENRYKELVSLGIKVQVLAIGNKAKLYFKRRPKFNVLKAFSLGQT-PSIKE
C. reinhardtii    GYNNFIIKKTEARYRELTAMGVKVNLVCVGRKGAQYFARRKQYNIVKSFSLGAA-PSTKE
V. carteri        GYNNFIIKKTEQRIRELTALGVKVNLVCVGRKGAQYFTKRKQYNVTKTFSLGAT-PSTKD
T. sociales       GYNNFIIKKSEKRIRELTALGLSVKVVCVGRKGGQYFNRRKQYNLVKSFILGAT-PTTKD
D. salina         GYNNFILKKTEQRFAELTALGLKVQVVAVGKKAQTYFKRRPKFNVIKEFGLGKS-PTTRD
C. eustigma       GYNNFILKKTEARFKELQALGLNVKVVAIGRKGAQYMKRRPKFNIVKEFSLGKA-PTMQD
H. oviformis      GFNNQSIKKAEARMKDLKELGVEFTVISVGKKGNSYFLRR-------NFSLGQT-PSTRD
                  *:**   **   *  ::   *:.  ::.:*.*.  :  :*        :      *: ::

A. thaliana 1     AQVIADDVFSLFVSEEVDKVELVYTKFVSLVKSDPVIHTLLPLSMKGESCDVKGECVDAI
A. thaliana 2     AQAVADDVFSLFISEEVDKVELLYTKFVSLVKSEPVIHTLLPLSPKGEICDINGTCVDAA
S. oleracea       AQAIADDVFSLFVSEEVDKVEMLYTKFVSLVKSDPVIHTLLPLSPKGEICDINGKCVDAA
C. braunii        AQAIADELFALFVSEEVDKVELLYTKFVSLVRSEVVIHTLLPLSPQGQVCDVDGNCVDAA
H. lacustris      AQAISDEVFSSFVSAEVDKVELVFTKFVSLISSTPTIQTLLPMTPAGELCNIDGTCVDAA
C. reinhardtii    AQGIADEIFASFIAQESDKVELVFTKFISLINSNPTIQTLLPMTPMGELCDVDGKCVDAA
V. carteri        AQGIADEIFASFIAQESDKVELVYTKFISLITSNPAIQTLLPMTPMGELCDVDGKCVDAA
T. sociales       AQAIADEIFASFIAQESDKVELVFTKFISLINSTPTIQTLLPMTTMGELCDVDGKCVDAA
D. salina         AQAISDTIFASFVSKEVDKVELVYTKFVSLVASTPTVQTVLPMAPAGELCNVDGTCVDAA
C. eustigma       SQAISDELFSSFVSEEVDKVELIYTKFVSLISSDPAVQTLLPLAPSGQLCNVDGTCVDAA
H. oviformis      AQGISDEILSSFVAAECDKVELLYTKFVSLITSNPTIQTLLPLTPTGELCGVDGNCVDAA
                  :* ::* ::: *:: * ****:::***:**: *  .::*:**::  *: *.:.* ****

A. thaliana 1     EDEMFRLTSKDGKLAVERTKLEVEKP-EISPLMQFEQDPVQILDAMMPLYLNSQILRALQ
A. thaliana 2     EDEFFRLTTKEGKLTVERETFRTPTA-DFSPILQFEQDPVQILDALLPLYLNSQILRALQ
S. oleracea       EDELFRLTTKEGKLTVERDMIKTETP-AFSPILEFEQDPAQILDALLPLYLNSQILRALQ
C. braunii        EDELFRLTTKEGKFVVEREVVRTSTE-SFEGILQFEQDPVQILDALLPLYMNSQVLRALQ
H. lacustris.     DDEIFKLTTRGGQFVVEREKAPIATD-ALDPSLIFEQEPSQILDALLPLYLNSCLLRSLQ
C. reinhardtii    DDEIFKLTTKGGEFAVEREKTTIETE-ALDPSLIFEQEPAQILDALLPLYMSSCLLRSLQ
V. carteri        DDEIFKLTTKGGEFAVEREKTTIATE-ALDPSLIFEQEPAQILDALLPLYMNSCLLRSLQ
T. sociales       DDEIFKLTTKGGAFAVERETTTITTE-ALDPSLIFEQEPAQILDALLPLYMNSCLLRSLQ
D. salina         NDEVFKLTSKGGQFSVERESSAIPTG-ELDPSLIFEQDPTQVLDALLPLYLNGCLLRSLQ
C. eustigma       NDEIFKLTSHNGKFEVTRESKPIDTDGGLDAGLIFEQEPAQILDSLLPLYLNAAMLRSLQ
H. oviformis      DDEVFRLTTKGGELSVEREKKPIETG-SLEPGLIFEQDPGQLLDALLPLYLNSCLLRALQ
                  :**.*:**:: * : * *      .   :.  : ***:* *:**:::***:.. :**:**
```

### Figure S7

```
A. thaliana 1     ESLASELASRMNAMSNATDNAVELKKNLTMAYNRARQAKITGELLEIVAGAEALRES
A. thaliana 2     ESLASELAARMSAMSSASDNASDLKKSLSMVYNRKRQAKITGEILEIVAGANAQV--
S. oleracea       ESLASELAARMTAMSNATDNANELKKTLSINYNRARQAKITGEILEIVAGANACV--
C. braunii        ESVASELASRMNAMNNASDNARDLKKNLSISYNRQRQAKITGEIMEIIAGANA----
H. lacustris      ESLASELAARMNAMGNATDNAKELRKVLTSSYNRKRQAKITQEISEITAGAAATS-G
C. reinhardtii    EALASELAARMNAMNNASDNAKELKKGLTVQYNKQRQAKITQELAEIVGGAAATS-G
V. carteri        EALASELAARMNAMNNASDNAKELRKALTVQYNKQRQAKITQELSEIVGGAAATS-G
T. sociales       EALASELAARMTAMNNASENAKELGKVLKVQYNKQRQAKITQELAEIVGGAAAAS-S
D. salina         ESLASELAARMNAMGTASDNAKELRKTLSNKYNKQRQARITQEISEIVSGASA----
C. eustigma       ESLASELAARMNAMANASDNAKELRKDLSAKYNRQRQAKITQELAELVGGAAATS--
H. oviformis      EALASELAARMNAMNSATDNA-ELIDGLTLSYNRARQAAITQEITEIVGGAAA----
                  *::*****:**.** .*::** :* . *.  **: *** ** *: *: .** *
```

# B

#### *Hyalomonas oviformis* δ subunit of $cF_OF_1$ ATPase

**TargetP 2.0 prediction**: Thylakoid luminal transfer peptide
CS pos: 33-34. VMA-KR. Pr: 0.5365
Chloroplast transfer peptide likelihood = 0.9146

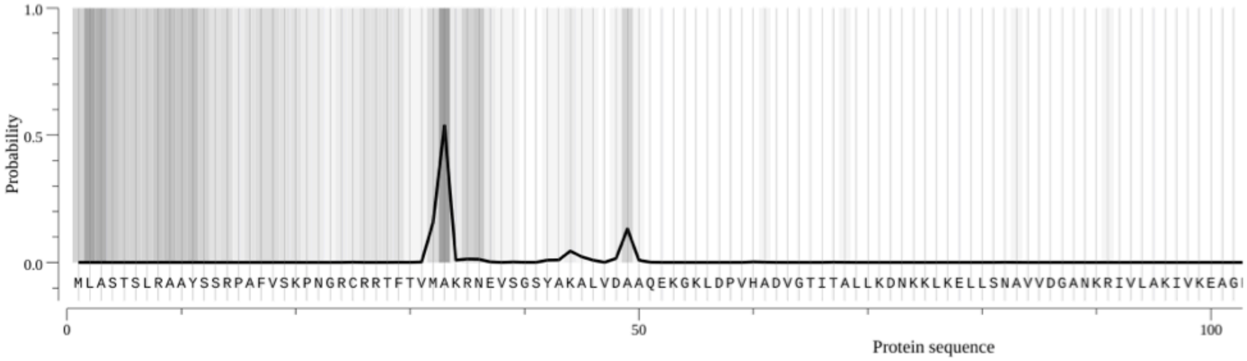


#### Multiple alignment δ subunit of $cF_OF_1$ ATPase

```
H. oviformis      ---------MLAST-------------------SLRAAYSSRPAFVSKPN---------
C. eustigma       ---------MIASK-------------------SLRAA-SSRAAGVSRVP---------
G. pectorale      ---------MLAAKS------------------SMR---ACKAS-TRAAP---------
V. carteri        ---------MLAAK-------------------SVVGQRAFKASATRAAP---------
C. reinhardtii    ---------MLAAK-------------------SIAGPRAFKASAVRAAP---------
T. sociales       ---------MLAAKS------------------SVMGQRVFKASAGRAAP---------
H. lacustris      MA-------ALTSKT------------------AMR---GLRASAPVRSI---------
D. salina.        M--------ALANQI------------------SRRGVSSTGKAPIARVPVARAP-IAS
A. thaliana       MASLQQTLFSLQSKL--------------PPSSFQIARSLPLRKTFPIRI----------
S. oleracea       MAALQNPV-ALQSRTTTAVAALSTSSTTSTPKPFSLSFSSSTATFNPLRLKILTASKLTA
                            :
```

### Figure S7

```
H. oviformis      GRCRRTFTVMAKRNEVSGSYAKALVDAAQEKGKLDPVHADVGTITALLKDNKKLKELLSN
C. eustigma       ARGRKAFTVLAKKNEVSESYAKALVDLADEKGKLEPVHADVDAVAALLKENQKLYELIAN
G. pectorale      RSARRVMVVMARKSEVSESYAKALVELADEKGKLEAVHADVDAVASLLKENKKLEALISN
V. carteri        KGARRAVVVMARKSEVSESYAKALVELADEKGKLEAVHADVDAVSSLMKENKKLSDLIMN
C. reinhardtii    KAGRRTVVVMARKNEVSESYAKALVELADEKGKLEAVHADVDAVAGLMKENAKLSALIMN
T. sociales       ---QRAVVAMARKNEVSESYAKALVELADEKGKLEAVHADVDAVASLMKENKKLAELIMN
H. lacustris      RSSRKAVCVQAKRNEVSESYAKALVELADEKGKLELVHADVDAISMLLQTNLKLTQLIMN
D. salina.        RSRRTAVKVQAVKNPVGESYAKALVDLANEKGKLEPIHADMDAVSQLMNTNKALSELVTN
A. thaliana       NNGGNAAGAR-MSATAASSYAMALADVAKRNDTMELTVTDIEKLEQVFSDP-QVLNFFAN
S. oleracea       KPRGGALGTR-MVDSTASRYASALADVADVTGTLEATNSDVEKLIRIFSEE-PVYYFFAN
                       .  .      ..  ** **.: *. ...::   :*:  :  ::.    :  :. *

H. oviformis      AVVDGANKRIVLAKIVKEAGFQQYTSNLLN------------------------------
C. eustigma       PVVDAEKKRSVLTKIGKEAGFHQYTMNFLNVLVQKDRLALLDEICESFEEQYCKLTDTQV
G. pectorale      PVVDGDKKKAVLAKVAKEAGFQQYTVNFLNLLVEKDRLSLVPEICEVFEELYCQLTDTQV
V. carteri        PIVDADKKRAVLAKIAKEAGFQQYTVNFLNLLVEKDRLSLVPEICECFEDLYCQMTDTQV
C. reinhardtii    PVVESDKKRAVLAKIAKEAGFQQYTINWLNLLVEKDRLSLVPEICECFEDLYCQMTDTQV
T. sociales       PVVDDDKKRAVLAKIGKEAGFQQYTVNFLNLLVAKDRLALVPEICETFEDLYCQMTDTQV
H. lacustris      PVVEADKKRAVLAKIGKEAGFNQYTNNFLNLLVQKDRLSLLEEICEEFEELYCKLTDTQ-
D. salina.        PVIDKEKKRAVLAKIGQEAGFQTYTNNFLNLLVEKDRMHLIYDICEAFEAQYCELTDTQV
A. thaliana       PTITVEKKRQVIDDIVKSSSLQSHTSNFLNVLVDANRINIVTEIVKEFELVYNKLTDTQL
S. oleracea       PVISIDNKRSVLDEIITTSGLQPHTANFINILIDSERINLVKEILNEFEDVFNKITGTEV
                  . :   :*: *: .:   :.:: :* * :*

H. oviformis      --------------------------------------SLIAGFVVEY---GTSQIDLSI
C. eustigma       AVVRSAVKLEQEQQFMIAKKLQELTGSKNIKLRPVIDQTLIAGFVVEY---GSSQIDLSI
G. pectorale      ATLRSAVKLEQEQQFLIAKKLQELTGSKNIKLKPVIDSSLIAGFVVEY---GSSQIDLSV
V. carteri        ATLRSAVKLEQEQQFLIAKKLQELTGSKNIKLKPVIDSSLIAGFVVEY---GSSQIDLSI
C. reinhardtii    ATLRSAVKLEQEQQFLIAKKLQELTGSKNIKLKPVIDSSLIAGFVVEY---GSSQIDLSV
T. sociales       ATLRSAVKLEQEQQFLIAKKLQELTGSKNIKLKPVIDSSLIAGFVVEY---GSSQIDLSV
H. lacustris      ------------------------------------------------------------
D. salina.        ALLRSAVKLEQEQQFLIAKKIQELSGSKNIKLKPTIDSSVIAGFIVEY---GSQQIDMSV
A. thaliana       AEVRSVVKLEAPQLAQIAKQVQKLTGAKNVRVKTVIDASLVAGFTIRYGESGSKLIDMSV
S. oleracea       AVVTSVVKLENDHLAQIAKGVQKITGAKNVRIKTVIDPSLVAGFTIRYGNEGSKLVDMSV

H. oviformis      KGQIDKVAEQLTSEMKLRLA
C. eustigma       KNQIEKVAEQLTKEMTLRMA
G. pectorale      RGQIEKVADQLTTDMTAKLA
V. carteri        RGQIERVADQLTKEMTAKLA
C. reinhardtii    RGQIERVADQLTKEMTAKLS
T. sociales       RGQIERVAEAVTKEMTAKLA
H. lacustris      --------------------
D. salina.        RGQVDNVTEELSKQMAMKMA
A. thaliana       KKQLEDIASQLELGEIQLAT
S. oleracea       KKQLEEIAAQLEMDDVTLAV
```

**Supplemental Figure S7.** Multiple protein sequence alignments of the **A**) γ and **B**) δ subunits of the cF$_0$-CF$_1$ ATPase from diverse viridiplants. Amino acid residues highlighted in yellow colour represent the transit peptides for import into the plastid predicted by TargetP 2.0 [http://www.cbs.dtu.dk/services/TargetP/] (Armenteros et al., 2019). In the case of the γ subunit multiple alignment (panel **A**), the columns emphasized in black background indicate the two cysteine residues typically involved in the thiol-modulation of the cF$_0$-CF$_1$ ATPase activity.

### Figure S8

#### Multiple sequence alignments of ChlB proteins from *Chlorogonium* species and *Hyalogonium fusiforme*

```
C.euchlorum      MKLAYWMYAGPAHIGTLRVASSFKNVHAIMHAPLGDDYFNVMRSMLERERDFTPVTASIV
C.capillatum     MKLAYWMYAGPAHIGTLRVASSFKNVHAIMHAPLGDDYFNVMRSMLERERDFTPVTASIV
H.fusiforme      MKLAYWMYAGPAHIGTLRVTSSFKNVHAIMHGPIGDSFFPVLSSMLSRSRDFAPVTTSIL
                 *******************:***********.*:**.:* *: ***.*.***:***:**:

C.euchlorum      DRHVLARGSQEKVVENFHRKEKEERPDLILLTPTCTSSILQEDLQNFVNRAK-----N-T
C.capillatum     DRHVLARGSQEKVVENFHRKEKEERPDLILLTPTCTSSILQEDLQNFVNRAN-----NEM
H.fusiforme      DRHVLARGSQEKVIENLRRKEKEEHPDLIVLTPTCTSSILQEDLQNFVIAAKQHIVQNDL
                 *************:**::******:****:******************  *:     *

C.euchlorum      SSNCDVLLADVNHYRVNEFQAADRTLEQIVRFYIEKAKKDNYYVPSLPLLADNPNQPLIS
C.capillatum     SSNCDVLLADVNHYRVNEFQAADRTLEQIVRFYIEKAKKDHFY----------NNEPFID
H.fusiforme      SSECDVLFIDSNHYRLNEHQAADLALEQIVRHYLEKAKKED-------------------
                 **:****: * ****:**.**** :******.*:*****:.

C.euchlorum      GEKVGVIPQGSALGNGGLEVGLEGESPFRTNNKQSLGLETQRLCEHQAKDLMDEQLWNNQ
C.capillatum     NYCVGEAEKGVFL-------------------KEKEG-----------------------
H.fusiforme      ------------------------------------------------------------

C.euchlorum      QQQIGSTSNTFVLNGICKTQKPSANILGVFTLGFHNLHDCRELKNLLTDLGIEVNEIIPE
C.capillatum     ------------VNGFCKTPKPSANILGVFTLGFHNLHDCRELKNLLTDLGIEVNEIIPE
H.fusiforme      ------------TGKSMRSPQPSCNILGMFTLGFHSQHDLRELKNLLNDLGVEVNQVIPE
                              .   :: :**.****:******. ** *******.***:***::***

C.euchlorum      GGTVNCLKNLPKAWFNIVPYRENGLMTAVYLEKEFSMPYTNIIPMGIVDTAAFIREIGNI
C.capillatum     GGTIKNLKNLPKAWFNIVPYRENGLMTAVYLEKEFSMPYTNITPMGIVDTAAFIREIGNI
H.fusiforme      GGSVTAIKNLRNAWFNIVPFRDAGLMTAVYLAKEFNMPYTTITPIGLIDTAAFAREISLI
                 **::. :*** :*******:*: ******** ***.****.* *:*::***** ***. *

C.euchlorum      CRSYPGQSPGLAAQMYNSDSSSSILNYMQSYPKTAPKEGSLQDYDKVLGTKPWEGTKPLI
C.capillatum     CRSINTQ-----------------ISDERRKTKMVGSKGDVHTQD--------SGRKPLI
H.fusiforme      CRS------------------------------AVASDSALLKST--------TGFSPLI
                 ***                               . ... :             * .***

C.euchlorum      LNQIACEKYIEKQTYFVSQAAWFSRSIDCQNLTGKKTVVFGDATHAASMTKILVREMGIH
C.capillatum     LNEIACEKYIEKQTYFVSQAAWFSRSIDCQNLTGKKTVVFGDATHAASMTKILVREMGIH
H.fusiforme      LNQEGLEKYIEKQTSFVSQAAWFNRSVDSNNLTGKRVFVFGSASHSASITKLLVREMGVH
                 **: . ******** ********.**:*.:*****:..***.*:*:**:**:******:*

C.euchlorum      VVCAGTYCKHDADWFREQVSGFCDQILITDDHSQIGDIIANILPAAIFGTQMERHVGKRL
C.capillatum     VVCAGTYCKHDADWFREQVSGFCDQILITDDHSQIGDIIAQIEPAAIFGTQMERHVGKRL
H.fusiforme      VVCAGTFQKHDADWFTHQVSGFCNKILITDDHSQVADVIANLEVDAIFGTQSERHVGKRL
                 ******: ******* .******::*********:.*:**::   ****** ********

C.euchlorum      DIPCGVISAPVHIQNFPLGFRPFLGYEGTNQISDLVYNSFSLGMEDHLLEIFNGHDNKEV
C.capillatum     DIPCGVISAPVHIQNFPLGFRPFLGYEGTNQISDLVYNSFSLGMEDHLLEIFNGHDNKEV
H.fusiforme      NLPCGVISAPVHIQNYPMGFKPFLGYEGANQIADLIYNSFALGMEEHMLDMFKGHDTAEI
                 ::*************:*:**:*******:***:**:****:****:*:*::*:***. *:

C.euchlorum      VTRSYSSE---NNILWTKEALAELSKIPGFVRAKVKRNTEKYAQTQRIQLITLEAMYAAK
C.capillatum     VTRSYSSE---NNILWTKEALAELEKVPGFVRAKVKRNTEKYAQTQKIQNITLEAMYAAK
H.fusiforme      TSRSYSSDASSTKLEWTKEALAELSKIPGFVRSKVKRNTEKYAQTKQINLITLEVIYDAK
                 .:*****:   .:: *********.*:*****:************::*: ****.:* **

C.euchlorum      EAVSSDSKS
C.capillatum     EAVS-----
H.fusiforme      EAVS-----
                 ****
```

Residues involved in the binding of the [4Fe-4S] cluster

Residues involved in ChlL binding

Residues involved in stabilizing the [4Fe-4S] cluster

**Supplemental Figure S8**. Multiple protein sequence alignments of the ChlB from *Chlorogonium* species and *Hyalogonium fusiforme*. Residues highlighted in magenta and cyan represent amino acids involved in the binding and stabilization of the [4Fe-4S] cluster of the protochlorophyllide oxidoreductase. The residues with green background indicate amino acids participating in the interaction of ChlB (binding) with ChlL (Bröcker et al., 2010).
