## Supplemental Figure S9 for "The plastomes of *Hyalomonas oviformis* and *Hyalogonium fusiforme* evolved dissimilar architecture after the loss of photosynthesis"

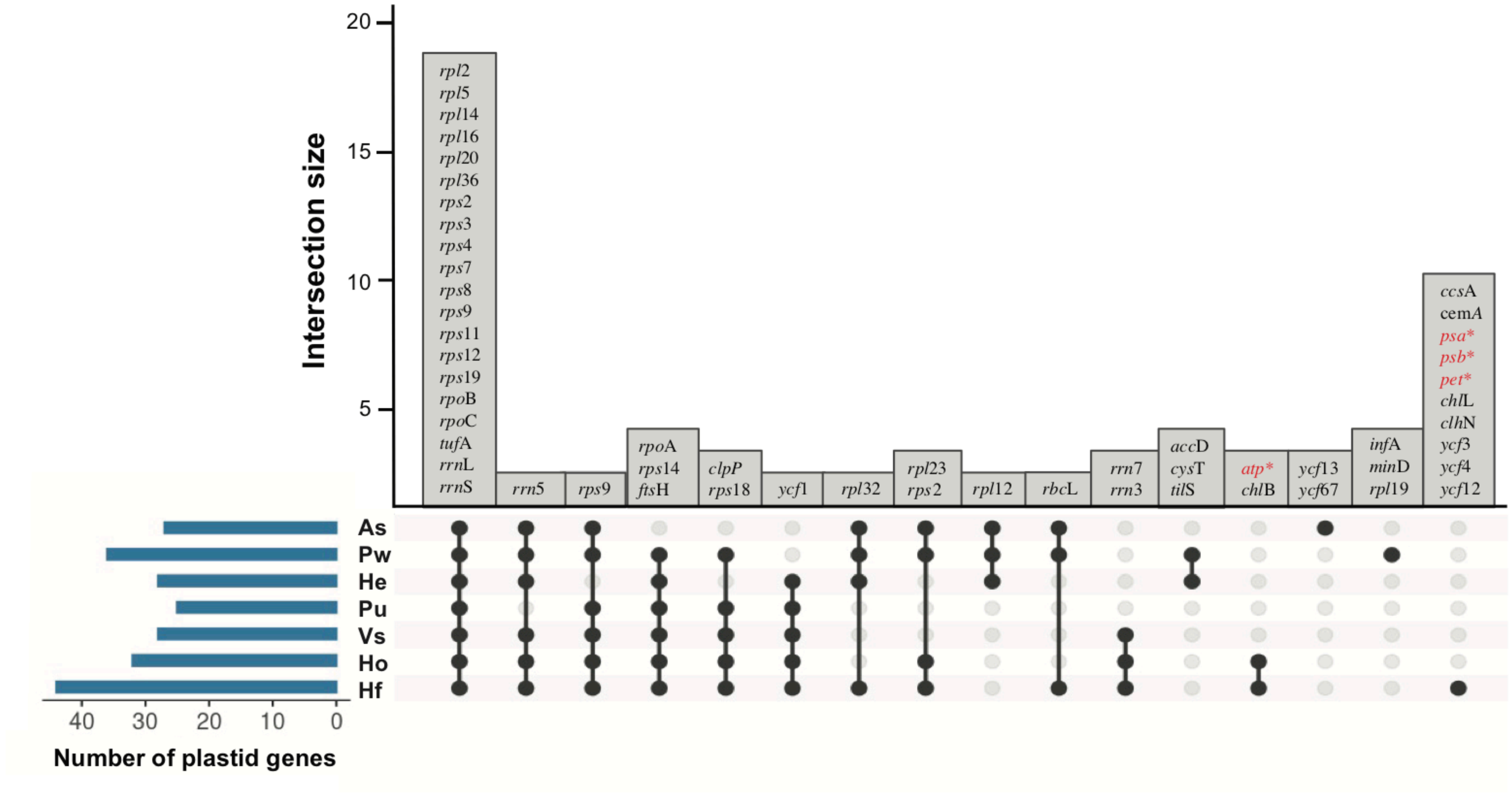


**Supplemental Figure S9**. Comparative content of protein-coding genes in plastid genomes of diverse non photosynthetic green algae and *Euglena* (*Astasia*) *longa*, which contains a secondary plastid of green algal origin. Gene names with asterisks refer to gene series encoding subunit of the CF-ATPase (*atp* series), the *b6f* complex (*pet* series), photosystem I (*psa* series) and photosystem II (*psb* series). Taxa codes: As, *Euglena longa*; Pw, *Prototheca wickerhamii*; He, *Helicosporidium* sp.; Pu, *Polytoma uvell*a; Vs, *Volvocales* NrCl902; Ho, *Hyalomonas oviformis*; Hf, *Hyalogonium fusiforme*.
