## Supplemental Table S1 for "The plastomes of *Hyalomonas oviformis* and *Hyalogonium fusiforme* evolved dissimilar architecture after the loss of photosynthesis"

**Table S1. RelTime estimated divergence times of non-photosynthetic chlamydomonadalean algae**

| | | Divergence times (MY) | | |
|---|---|---|---|---|
| **Species (strain)** | **Data** | **Mean** | **Minimum** | **Maximum** |
| *Hyalogonium fusiforme* (SAG 62-1c) | 16S rRNA + 18S rRNA | 77.1 | 33.1 | 179.4 |
| | 5 plastid proteins[1] | 45.4 | 23.1 | 89.3 |
| *Hyalomonas oviformis* (SAG 62-27) | 16S rRNA + 18S rRNA | 114.2 | 52..9 | 246.2 |
| | 5 plastid proteins[1] | 104.3 | 64.3 | 169.3 |
| *Polytoma uvella* (UTEX 964) | 16S rRNA + 18S rRNA | 121.7 | 58.0 | 255.3 |
| | 5 plastid proteins[1] | 92.0 | 48.0 | 176.2 |

1. Conceptual translation of the plastid genes *tuf*A, *rps*4, *rps*7, *rpl*2 and *rpl*5
Approximated divergence times were estimated with the RelTime approach

**Table S2. General information of the Illumina and PacBio assemblies**

| Species (Strain) | Sequencing technology | No. raw reads | Assembler | Assembly type | No. contigs | Largest fragment |
|---|---|---|---|---|---|---|
| *Hyalomonas oviformis* (SAG 62-27) | Illumina | 174,274,557 | SPAdes | Short read | 290,515 | 11,657,939 |
| *Hyalomonas chlamydogama* (SAG 11-48b) | Illumina | 76,681,410 | Ray | Short read | 115,721 | 451,573 |
| *Chlorogonium euchlorum* (SAG 12-2a) | Illumina | 175,527,406 | Ray | Short read | 161,821 | 114,504 |
| | | | SPAdes | Hybrid | 261,062 | 147,504 |
| | PacBio | 555,390 | Canu | Long read | 2,119 | 35,343 |
| *Hyalogonium fusiforme* (SAG 62-1c) | Illumina | 228,850,133 | Ray | Short read | 3,823,594 | 1,834,395 |
| | PacBio | 413,219 | Unicycler | Hybrid | 4,741 | 2,289,232 |
| | | | SPAdes | Hybrid | 189,424 | 1,954,482 |
| | | | Canu | Long read | 1,970 | 47,233 |

**Table S3**. Average read coverage estimated per genomic compartment

| **Average coverage (reads per nucleotide ± SD)** | ***Hyalomonas oviformis*** | ***Hyalomonas chlamydogama*** | ***Hyalogonium fusiforme*** | ***Chlorogonium euchlorum*** |
|---|---|---|---|---|
| Plastid protein-coding genes | 281 ± 91.8 | 5,030 ± 408.3 | 666 ± 118.2 | 17,377 ± 6,093.5 |
| GC % | 25.2 | 29.2 | 36.7 | 34.4 |
| rRNAs | 429 ± 431.8 | 9,504 ± 476.5 | 2,092 ± 3,441.6 | 37,942 ± 5,512.6 |
| Plastid scaffolds/ complete genome | 308 ± 155.8 | 5,011 ± 449.4 | 703.0 ± 631.7 | 16,521 ± 8,004.2 |
| GC % | 28.1 | 27.6 | 44.4 | 36.9 |
| Mitochondrial scaffolds / complete genome | 1,426 ± 73.5 | 4,300 ± 657 | 2,505 ± 482.2 | 34,545 ± 3,015 |
| GC % | 32.3 | 31.9 | 36.8 | 35.2 |
| Nuclear sequences* | 16.3 ± 28.9 | 43 ± 8.8 | 264 ± 850.4 | 262 ± 257.8 |
| GC % | 57.3 | 55.6 | 58.1 | 54.2 |

*Nuclear contigs identified by BLAST containing protein-coding genes typically present as single-copy in *Chlamydomonas reinhardtii*. The 15 genes considered encode heat shock protein 90A [HSP90A], heat shock protein 70 [HSP70], actin, tubulin beta-1 chain, tubulin alpha-1 chain, elongation factor 2 [EFG2], 20S proteasome beta subunit, 14-3-3 protein, elongation factor 1 alpha 1 [EF1A], RubV-2 helicase [reptin], AAA family ATPase [transitional endoplasmic reticulum], V-ATPase B subunit, ATP-dependent 26S proteasome regulatory subunit [RPT4], translation initiation factor 2 gamma [EIF2G], eukaryotic translation initiation factor 5A [EIF5A].

### Table S4. Comparative gene content of plastid genomes from diverse chlorophytes

| Category | Gene | CHL |  |  |  |  |  |  |  |  |  |  | TRE |  |
|---|---|---|---|---|---|---|---|---|---|---|---|---|---|---|
|  |  | *Ho* | *Hc* | *Hf* | *Cge* | *Cgc* | *Pu* | *Cl* | *Cr* | *Vs* | *Hl* | *Lp* | *He* | *Pw* |
| Ribosomal large subunit | *rpl1* |  |  |  |  |  |  |  |  |  |  |  |  |  |
|  | *rpl2* | ■ | ■ | ■ | ■ | ■ | ■ | ■ | ■ | ■ | ■ | ■ | ■ | ■ |
|  | *rpl3* |  |  |  |  |  |  |  |  |  |  |  |  |  |
|  | *rpl4* |  |  |  |  |  |  |  |  |  |  |  |  |  |
|  | *rpl5* | ■ | ■ | ■ | ■ | ■ | ■ | ■ | ■ | ■ | ■ | ■ | ■ | ■ |
|  | *rpl6* |  |  |  |  |  |  |  |  |  |  |  |  |  |
|  | *rpl11* |  |  |  |  |  |  |  |  |  |  |  |  |  |
|  | *rpl12* |  |  |  |  |  |  |  |  |  |  |  | ■ | ■ |
|  | *rpl13* |  |  |  |  |  |  |  |  |  |  |  |  |  |
|  | *rpl14* | ■ | ■ | ■ | ■ | ■ | ■ | ■ | ■ | ■ | ■ | ■ | ■ | ■ |
|  | *rpl16* | ■ | ■ | ■ | ■ | ■ | ■ | ■ | ■ | ■ | ■ | ■ | ■ | ■ |
|  | *rpl19* |  |  |  |  |  |  |  |  |  |  |  |  | ■ |
|  | *rpl20* | ■ | ■ | ■ | ■ | ■ | ■ | ■ | ■ | ■ | ■ | ■ | ■ | ■ |
|  | *rpl21* |  |  |  |  |  |  |  |  |  |  |  |  |  |
|  | *rpl22* |  |  |  |  |  |  |  |  |  |  |  |  |  |
|  | *rpl23* | ■ | ■ | ■ | ■ | ■ |  | ■ | ■ |  | ■ | ■ |  | ■ |
|  | *rpl24* |  |  |  |  |  |  |  |  |  |  |  |  |  |
|  | *rpl27* |  |  |  |  |  |  |  |  |  |  |  |  |  |
|  | *rpl29* |  |  |  |  |  |  |  |  |  |  |  |  |  |
|  | *rpl31* |  |  |  |  |  |  |  |  |  |  |  |  |  |
|  | *rpl32* |  |  | ■ | ■ | ■ |  |  |  |  | ■ | ■ | ■ | ■ |
|  | *rpl33* |  |  |  |  |  |  |  |  |  |  |  |  |  |
|  | *rpl34* |  |  |  |  |  |  |  |  |  |  |  |  |  |
|  | *rpl35* |  |  |  |  |  |  |  |  |  |  |  |  |  |
|  | *rpl36* | ■ | ■ | ■ | ■ | ■ | ■ | ■ | ■ | ■ | ■ | ■ | ■ | ■ |
| Ribosomal small subunit | *rps2* | ■ | ■ | ■ | ■ | ■ |  | ■ | ■ | ■ | ■ | ■ |  | ■ |
|  | *rps3* | ■ | ■ | ■ | ■ | ■ | ■ | ■ | ■ | ■ | ■ | ■ | ■ | ■ |
|  | *rps4* | ■ | ■ | ■ | ■ | ■ | ■ | ■ | ■ | ■ | ■ | ■ | ■ | ■ |
|  | *rps5* |  |  |  |  |  |  |  |  |  |  |  |  |  |
|  | *rps6* |  |  |  |  |  |  |  |  |  |  |  |  |  |
|  | *rps7* | ■ | ■ | ■ | ■ | ■ | ■ | ■ | ■ | ■ | ■ | ■ | ■ | ■ |
|  | *rps8* | ■ | ■ | ■ | ■ | ■ | ■ | ■ | ■ | ■ | ■ | ■ | ■ | ■ |
|  | *rps9* | ■ | ■ | ■ | ■ | ■ | ■ | ■ | ■ | ■ | ■ | ■ |  | ■ |
|  | *rps10* |  |  |  |  |  |  | ■ | ■ |  | ■ | ■ |  |  |
|  | *rps11* | ■ | ■ | ■ | ■ | ■ | ■ | ■ | ■ | ■ | ■ | ■ | ■ | ■ |
|  | *rps12* |  | ■ | ■ | ■ | ■ | ■ | ■ | ■ |  | ■ | ■ | ■ | ■ |
|  | *rps13* |  |  |  |  |  |  |  |  |  |  |  |  |  |
|  | *rps14* | ■ | ■ | ■ | ■ | ■ | ■ | ■ | ■ | ■ | ■ | ■ | ■ | ■ |
|  | *rps16* |  |  |  |  |  |  |  |  |  |  |  |  |  |
|  | *rps17* |  |  |  |  |  |  |  |  |  |  |  |  |  |
|  | *rps18* | ■ | ■ | ■ | ■ | ■ | ■ | ■ | ■ | ■ | ■ | ■ |  | ■ |
|  | *rps19* | ■ | ■ | ■ | ■ | ■ | ■ | ■ | ■ | ■ | ■ | ■ | ■ | ■ |
|  | *rps20* |  |  |  |  |  |  |  |  |  |  |  |  |  |
| Transcription/translation | *rpoA* | ■ | ■ | ■ | ■ | ■ | ■ | ■ | ■ | ■ | ■ | ■ | ■ | ■ |
|  | *rpoB1* | ■ | ■ | ■ | ■ | ■ | ■ | ■ | ■ | ■ | ■ | ■ | ■ | ■ |
|  | *rpoB2* | ■ | ■ |  | ■ | ■ | ■ | ■ | ■ | ■ | ■ | ■ |  |  |
|  | *rpoC1* | ■ | ■ | ■ | ■ | ■ | ■ | ■ | ■ | ■ | ■ | ■ | ■ | ■ |
|  | *rpoC2* | ■ | ■ | ■ | ■ | ■ | ■ | ■ | ■ | ■ | ■ | ■ | ■ | ■ |
|  | *infB* |  |  |  |  |  |  |  |  |  |  |  |  | ■ |
|  | *infC* |  |  |  |  |  |  |  |  |  |  |  |  |  |
|  | *tsf* |  |  |  |  |  |  |  |  |  |  |  |  |  |
|  | *tufA* | ■ | ■ | ■ | ■ | ■ | ■ | ■ | ■ | ■ | ■ | ■ | ■ | ■ |
| ATP synthase | *atpA* | ■ | ■ | ■ | ■ |  |  | ■ | ■ |  | ■ | ■ |  | ■ |
|  | *atpB* | ■ | ■ | ■ | ■ | ■ |  | ■ | ■ |  | ■ | ■ |  | ■ |
|  | *atpE* | ■ | ■ | ■ | ■ | ■ |  | ■ | ■ |  | ■ | ■ |  | ■ |
|  | *atpF* | ■ | ■ | ■ | ■ | ■ |  | ■ | ■ |  | ■ | ■ |  | ■ |
|  | *atpH* | ■ | ■ | ■ | ■ | ■ |  | ■ | ■ |  | ■ | ■ |  | ■ |
|  | *atpI* | ■ | ■ | ■ | ■ | ■ |  | ■ | ■ |  | ■ | ■ |  | ■ |
| Miscellaneous | *ccsA* |  | ■ | ■ | ■ | ■ |  | ■ | ■ |  | ■ |  |  |  |
|  | *cemA* |  | ■ | ■ | ■ | ■ |  | ■ | ■ |  | ■ |  |  |  |
|  | *clpP* | ■ | ■ | ■ | ■ | ■ | ■ | ■ | ■ | ■ | ■ | ■ |  |  |
| chlorophyll biosynthesis | *chlB* | ■ | ■ | ■ | ■ | ■ |  | ■ | ■ |  |  |  |  |  |
|  | *chlL* |  | ■ | ■ | ■ | ■ |  | ■ | ■ |  |  |  |  |  |
|  | *chlN* |  | ■ | ■ | ■ |  |  | ■ | ■ |  |  |  |  |  |
| cytochrome b6/f | *petA* |  | ■ | ■ | ■ | ■ |  | ■ | ■ |  | ■ |  |  |  |
|  | *petB* |  | ■ | ■ | ■ | ■ |  | ■ | ■ |  | ■ |  |  |  |
|  | *petD* |  | ■ | ■ | ■ | ■ |  | ■ | ■ |  | ■ |  |  |  |
|  | *petG* |  | ■ | ■ | ■ | ■ |  | ■ | ■ |  | ■ |  |  |  |
|  | *petL* |  | ■ | ■ | ■ | ■ |  | ■ | ■ |  | ■ |  |  |  |
| Photosystem I | *psaB* |  | ■ | ■ | ■ | ■ |  | ■ | ■ |  | ■ |  |  |  |
|  | *psaC* |  |  |  |  |  |  | ■ | ■ |  | ■ |  |  |  |
|  | *psaI* |  | ■ | ■ | ■ | ■ |  |  |  |  |  |  |  |  |
|  | *psaJ* |  |  |  |  |  |  | ■ | ■ |  | ■ |  |  |  |
|  | *psaM* |  | ■ | ■ | ■ | ■ |  |  |  |  |  |  |  |  |
|  | *psaA* |  | ■ | ■ | ■ | ■ |  | ■ | ■ |  | ■ |  |  |  |
| Photosystem II | *psbA* |  | ■ | ■ | ■ | ■ |  | ■ | ■ |  | ■ |  |  |  |
|  | *psbB* |  | ■ | ■ | ■ | ■ |  | ■ | ■ |  | ■ |  |  |  |
|  | *psbC* |  | ■ | ■ | ■ | ■ |  | ■ | ■ |  | ■ |  |  |  |
|  | *psbD* |  | ■ | ■ | ■ | ■ |  | ■ | ■ |  | ■ |  |  |  |
|  | *psbE* |  | ■ | ■ | ■ | ■ |  | ■ | ■ |  | ■ |  |  |  |
|  | *psbF* |  | ■ | ■ | ■ | ■ |  | ■ | ■ |  | ■ |  |  |  |
|  | *psbH* |  | ■ | ■ | ■ | ■ |  | ■ | ■ |  | ■ |  |  |  |
|  | *psbI* |  | ■ | ■ | ■ | ■ |  | ■ | ■ |  | ■ |  |  |  |
|  | *psbJ* |  | ■ | ■ | ■ | ■ |  | ■ | ■ |  | ■ |  |  |  |
|  | *psbK* |  | ■ | ■ | ■ | ■ |  | ■ | ■ |  | ■ |  |  |  |
|  | *psbL* |  | ■ | ■ | ■ | ■ |  | ■ | ■ |  | ■ |  |  |  |
|  | *psbM* |  | ■ | ■ | ■ | ■ |  | ■ | ■ |  | ■ |  |  |  |
|  | *psbN* |  | ■ | ■ | ■ | ■ |  | ■ | ■ |  | ■ |  |  |  |
|  | *psbT* |  | ■ | ■ | ■ | ■ |  | ■ | ■ |  | ■ |  |  |  |
|  | *psbZ* |  | ■ | ■ | ■ | ■ |  | ■ | ■ |  | ■ |  |  |  |
|  | *psb30* |  |  |  |  |  |  | ■ | ■ |  |  |  |  |  |
| Rubisco | *rbcL* |  | ■ | ■ | ■ | ■ |  | ■ | ■ |  | ■ |  |  | ■ |
| Conserved genes | *ycf1* | ■ | ■ | ■ | ■ | ■ | ■ | ■ | ■ | ■ | ■ | ■ |  |  |
|  | *ycf3* |  | ■ | ■ | ■ | ■ |  | ■ | ■ |  | ■ |  |  |  |
|  | *ycf4* |  | ■ | ■ | ■ | ■ |  | ■ | ■ |  | ■ |  |  |  |
|  | *ycf12* |  | ■ | ■ | ■ | ■ |  | ■ | ■ |  | ■ |  |  |  |
|  | *ftsH* | ■ | ■ | ■ | ■ | ■ | ■ | ■ | ■ | ■ | ■ | ■ |  |  |

| Category | Gene | CHL |  |  |  |  |  |  |  |  |  |  | TRE |  |
|---|---|---|---|---|---|---|---|---|---|---|---|---|---|---|
|  |  | *Ho* | *Hc* | *Hf* | *Cge* | *Cgc* | *Pu* | *Cl* | *Cr* | *Vs* | *Hl* | *Lp* | *He* | *Pw* |
| Ribosomal RNAs | *rrnL* | ■ | ■ | ■ | ■ | ■ | ■ | ■ | ■ | ■ | ■ | ■ |  |  |
|  | *rrnS* | ■ | ■ | ■ | ■ | ■ | ■ | ■ | ■ | ■ | ■ | ■ |  |  |
|  | *rrn5* | ■ | ■ |  | ■ | ■ |  | ■ | ■ | ■ | ■ |  |  |  |
|  | *rrn7* | ■ |  |  | ■ |  |  | ■ | ■ | ■ | ■ |  |  |  |
|  | *rrn3* | ■ |  |  | ■ |  |  | ■ | ■ | ■ | ■ |  |  |  |
| Transfer RNAs | trnA (UGC) | ■ | ■ | ■ | ■ | ■ | ■ | ■ | ■ | ■ | ■ | ? | ■ | ■ |
|  | trnA (UGC) |  | ■ |  | ■ |  |  | ■ | ■ |  | ■ | ? |  |  |
|  | trnC (GCA) | ■ | ■ | ■ | ■ | ■ | ■ | ■ | ■ | ■ | ■ | ? | ■ | ■ |
|  | trnD (GUC) | ■ | ■ | ■ | ■ | ■ | ■ | ■ | ■ | ■ | ■ | ? | ■ | ■ |
|  | trnE (UUC) | ■ | ■ |  | ■ | ■ | ■ | ■ | ■ | ■ | ■ | ? | ■ | ■ |
|  | trnE (UUC) | ■ | ■ |  | ■ | ■ |  | ■ | ■ | ■ | ■ | ? |  |  |
|  | trnF (GAA) | ■ | ■ | ■ | ■ | ■ |  | ■ | ■ | ■ | ■ | ? | ■ | ■ |
|  | trnG (GCC) | ■ | ■ | ■ | ■ | ■ | ■ | ■ | ■ | ■ | ■ | ? | ■ | ■ |
|  | trnG (UCC) | ■ | ■ |  | ■ | ■ | ■ | ■ | ■ | ■ | ■ | ? |  | ■ |
|  | trnH (GUG) | ■ | ■ | ■ | ■ | ■ | ■ | ■ | ■ | ■ | ■ | ? | ■ | ■ |
|  | trnI (GAU) | ■ | ■ | ■ | ■ | ■ | ■ | ■ | ■ | ■ | ■ | ? | ■ | ■ |
|  | trnI (GAU) |  | ■ |  | ■ | ■ |  | ■ | ■ |  | ■ | ? | ■ |  |
|  | trnI (GAU) |  |  |  | ■ |  |  |  |  |  |  | ? |  |  |
|  | trnI (CAU) |  |  |  | ■ |  |  |  |  |  | ■ | ? |  |  |
|  | trnK (UUU) | ■ | ■ |  | ■ | ■ | ■ | ■ | ■ | ■ | ■ | ? | ■ | ■ |
|  | trnK (UUU) |  |  |  |  |  | ■ |  |  |  |  | ? |  |  |
|  | trnL(CAA) | ■ | ■ | ■ | ■ | ■ |  | ■ |  |  |  | ? | ■ | ■ |
|  | trnL(CAA) | ■ | ■ | ■ | ■ | ■ |  | ■ |  |  |  | ? | ■ | ■ |
|  | trnL (UAA) |  |  |  |  |  | ■ |  | ■ | ■ | ■ | ? |  |  |
|  | trnL (UAG) |  |  |  |  |  |  |  | ■ | ■ | ■ | ? |  |  |
|  | trnM (CAU) | ■ | ■ | ■ | ■ | ■ | ■ | ■ | ■ | ■ | ■ | ? | ■ | ■ |
|  | trnM (CAU) | ■ | ■ |  | ■ | ■ | ■ | ■ | ■ | ■ | ■ | ? | ■ | ■ |
|  | trnM (CAU) | ■ | ■ |  | ■ |  | ■ |  | ■ | ■ |  | ? |  | ■ |
|  | trnN (GUU) | ■ | ■ | ■ | ■ | ■ | ■ | ■ | ■ | ■ | ■ | ? | ■ | ■ |
|  | trnP (UGG) | ■ | ■ | ■ | ■ | ■ | ■ | ■ | ■ | ■ | ■ | ? | ■ | ■ |
|  | trnQ (UUG) | ■ | ■ | ■ | ■ | ■ | ■ | ■ | ■ | ■ | ■ | ? | ■ | ■ |
|  | trnQ (UUG) |  |  |  |  |  | ■ |  |  |  |  | ? |  |  |
|  | trnR (ACG) | ■ | ■ | ■ | ■ | ■ | ■ | ■ | ■ | ■ | ■ | ? | ■ | ■ |
|  | trnR (AGA) | ■ | ■ | ■ | ■ | ■ |  | ■ | ■ |  |  | ? | ■ | ■ |
|  | trnR(UCG) |  |  |  |  |  | ■ |  |  |  |  | ? |  | ■ |
|  | trnR(CCG) |  |  |  |  |  |  |  |  |  |  | ? |  |  |
|  | trnR(CCU) |  |  |  |  |  |  |  |  |  | ■ | ? |  |  |
|  | trnR(UCU) |  |  |  |  |  | ■ |  |  | ■ | ■ | ? |  |  |
|  | trnS (GCU) | ■ | ■ | ■ | ■ | ■ | ■ | ■ | ■ | ■ | ■ | ? | ■ | ■ |
|  | trnS (GCU) |  |  |  |  |  |  |  |  | ■ |  | ? |  |  |
|  | trnS (GGA) | ■ | ■ |  | ■ | ■ |  | ■ |  |  | ■ | ? | ■ | ■ |
|  | trnS (UGA) |  |  |  |  |  | ■ |  | ■ |  | ■ | ? |  |  |
|  | trnT (UGU) | ■ | ■ | ■ | ■ | ■ | ■ | ■ | ■ | ■ | ■ | ? | ■ | ■ |
|  | trnV (UAC) | ■ | ■ | ■ | ■ | ■ | ■ | ■ | ■ | ■ | ■ | ? | ■ | ■ |
|  | trnV (UAC) |  |  | ■ |  |  |  |  |  | ■ |  | ? |  |  |
|  | trnW (CCA) | ■ | ■ |  | ■ | ■ | ■ | ■ | ■ |  | ■ | ? | ■ | ■ |
|  | trnW (ACA) |  |  |  |  |  |  |  |  | ■ |  | ? |  |  |
|  | trnW (UCA) |  |  |  |  |  |  |  |  | ■ |  | ? |  |  |
|  | trnW (UCA) |  |  |  |  |  |  |  |  | ■ |  | ? |  |  |
|  | trnY (GUA) | ■ | ■ | ■ | ■ | ■ | ■ | ■ | ■ | ■ | ■ | ? | ■ | ■ |
|  | trnY (GUA) |  |  | ■ |  |  |  |  |  |  |  | ? |  |  |

**Gray rows/cells indicate nonphotosynthetic taxa**

**CLH, Chlamydomonadaceae**
***Ho***, *Hyalomanas oviformis*
***Hc***, *Hyalomonas chlamydogama*
***Hf***, *Hyalogonium fusiforme*,
***Cge***, *Chlorogonium euchlorum*
***Cgc***, *Chlorogonium capillatum*
***Pu***, *Polytoma uvella*
***Cl***, *Chlamydomonas leiostraca*
***Cr***, *Chlamydomonas reinhardtii*
***Vs***, *Volvocales sp. NrCl902*
***Hl***, *Hafniomonas laevis*
***Lp***, *Leontynka pallida*

**TRE, Trebouxiophyceae**
***He***, *Helicosporidium sp.*
***Pw***, *Prototheca wickerhamii*

**Table S5.** Size of predicted plastid protein-coding regions in *Hyalogonium fusiforme* and *Chlorogonium euchlorum*.

| ***Hyalogonium fusiforme*** SAG 62-1c | | ***Chlorogonium euchlorum*** SAG 12-2a | |
|---|---|---|---|
| **Gene or coding region** | **Coding region size (bp)** | **Gene or coding region** | **Coding region size (bp)** |
| *atp*A 1[2] | 37 | *atp*A | 1521 |
| *atp*A 2[2] | 687 | | |
| *atp*A 3[2] | 360 | | |
| *atp*B 1[2] | 216 | *atp*B | 1452 |
| *atp*B 2[2] | 563 | | |
| *atp*B 3[2] | 202 | | |
| *atp*B 4[2] | 393 | | |
| *atp*E | 405 | *atp*E | 408 |
| *atp*F | 534 | *atp*F1 | 453 |
| | | *atp*F2 | 621 |
| *atp*H | 249 | *atp*H | 249 |
| *atp*I | 741 | *atp*I | 720 |
| *ccs*A | 1569 | *ccs*A | 1536 |
| *cem*A | 858 | *cem*A | 1626 |
| *chl*B | 1608 | *chl*B | 1983 |
| *chl*L | 912 | *chl*L | 921 |
| *chl*N | 1377 | *chl*N | 1590 |
| *clp*P | 1743 | *clp*P | 1938 |
| *fts*H | 11319 | *fts*H | 11391 |
| *pet*A | 945 | *pet*A | 945 |
| *pet*B | 648 | *pet*B | 648 |
| *pet*D | 483 | *pet*D | 483 |
| *pet*G | 78 | *pet*G | 114 |
| *pet*L | 99 | *pet*L | 99 |
| *psa*A 1[2] | 138 | *psa*A 1[2] | 141 |
| *psa*A 2[2] | 141 | *psa*A 2[2] | 180 |
| *psa*A 3[2] | 1986 | *psa*A 3[2] | 1986 |
| *psa*B | 2208 | *psa*B | 2208 |
| *psa*C | 246 | *psa*C | 246 |
| *psa*J | 111 | *psa*J | 126 |
| *psb*A exon 1[1] | 553 | *psb*A exon 1[1] | 276 |
| *psb*A exon 2[1] | 440 | *psb*A exon 2[1] | 138 |

| | | | |
|---|---|---|---|
| *psb*A exon 3[1] | 63 | *psb*A exon 3[1] | 134 |
| | | *psb*A exon 4[1] | 242 |
| | | *psb*A exon 5[1] | 263 |
| *psb*B 1[2] | 405 | *psb*B | 1527 |
| *psb*B 2[2] | 1191 | | |
| *psb*C 1[2] | 7773 | *psb*C exon 1[1] | 720 |
| *psb*C 2[2] | 399 | *psb*C exon 2[1] | 504 |
| *psb*C 3[2] | 204 | | |
| *psb*C 4[2] | 29 | | |
| *psb*C 5[2] | 31 | | |
| *psb*D1[2] | 954 | *psb*D exon 1[1] | 545 |
| | | *psb*D exon 2[1] | 195 |
| | | *psb*D exon 3[1] | 319 |
| *psb*D2[2] | 171 | | |
| *psb*E1 | 105 | *psb*E | 252 |
| *psb*E2 | 159 | | |
| *psb*F1 | 94 | *psb*F | 135 |
| *psb*F2 | 41 | | |
| *psb*H | 264 | *psb*H | 264 |
| *psb*I1[2] | 141 | *psb*I | 114 |
| *psb*I2[2] | 30 | | |
| *psb*J | 138 | *psb*J | 129 |
| *psb*K1[2] | 51 | *psb*K | 141 |
| *psb*K2[2] | 90 | | |
| *psb*L | 117 | *psb*L | 117 |
| *psb*M | 105 | *psb*M | 108 |
| *psb*N | 129 | *psb*N | 135 |
| *psb*T1[2] | 135 | *psb*T | 96 |
| *psb*T2[2] | 21 | | |
| *psb*Z1[2] | 106 | *psb*Z | 189 |
| *psb*Z2[2] | 138 | | |
| *rbc*L 1[2] | 1014 | *rbc*L exon 1[1] | 699 |
| *rbc*L 2[2] | 384 | *rbc*L exon 2[1] | 729 |
| *rpl*14 | 360 | *rpl*14 | 360 |
| *rpl*16 | 438 | *rpl*16 | 426 |
| *rpl*2 | 828 | *rpl*2 | 828 |
| *rpl*20 | 339 | *rpl*20 | 339 |

| | | | |
|---|---|---|---|
| *rpl*23 | 375 | *rpl*23 | 390 |
| *rpl*32 | 177 | *rpl*32 1[2] | 13 |
| | | *rpl*32 2[2] | 195 |
| *rpl*36 | 114 | *rpl*36 | 114 |
| *rpl*5 | 543 | *rpl*5 | 543 |
| *rpo*A | 1452 | *rpo*A | 2250 |
| *rpo*B1 | 2490 | *rpo*Ba | 2595 |
| *rpo*B2 | 4329 | *rpo*Bb | 4155 |
| *rpo*C1 | 5769 | *rpo*C1 | 5400 |
| *rpo*C2 | 10473 | *rpo*C2 | 9117 |
| *rps*11 | 393 | *rps*11 | 393 |
| *rps*12 | 417 | *rps*12 | 402 |
| *rps*14 | 300 | *rps*14 | 303 |
| *rps*18 | 498 | *rps*18 | 492 |
| *rps*19 | 279 | *rps*19 | 324 |
| *rps*2 | 1818 | *rps*2 | 1746 |
| *rps*3 | 2403 | *rps*3 | 2307 |
| *rps*4 | 792 | *rps*4 | 816 |
| *rps*7 | 507 | *rps*7 | 507 |
| *rps*8 | 435 | *rps*8 | 435 |
| *rps*9 | 759 | *rps*9 | 540 |
| *tuf*A | 1257 | *tuf*A | 1257 |
| *ycf*1 | 7557 | *ycf*1 | 7371 |
| *ycf*12 | 93 | *ycf*12 | 102 |
| *ycf*3 | 513 | *ycf*3 | 525 |
| *ycf*4 1[2] | 255 | *ycf*4 | 630 |
| *ycf*4 2[2] | 384 | | |
| *ycf*4 3[2] | 237 | | |

[1]Exon/introns predicted with RNAweasel (https://megasun.bch.umontreal.ca/RNAweasel/; Lang et al. 2007)
[2]Identified fragments of typical plastid protein-coding genes not predicted as exons.

**Lang BF, Laforest MJ, Burger G** (2007) Mitochondrial introns: a critical view. Trends Genet **23**:119–125.

**Table S6** Functional classification of the *Hyalogonium fusiforme* TrRnEn-like ORFs using BLAST and HMMER. (Altschul *et al.*, 1990; Finn *et al.*, 2011).

| Endonuclease | Length (bp) | BLAST annotation | HMMER domains | E-value | Description |
|---|---|---|---|---|---|
| ORF 1 | 1788 | reverse transcriptase, II intron maturase | RVT_1 | 6.70E-21 | Reverse transcriptase (RNA-dependent DNA polymerase) |
| | | | RVT_N | 8.40E-28 | N-terminal domain of reverse transcriptase |
| | | | GIIM | 5.80E-22 | Group II intron, maturase-specific domain |
| ORF 2 | 873 | reverse transcriptase/ group II intron maturase / HNH | GIIM | 2.50E-18 | Group II intron, maturase-specific domain |
| | | | HNH | 2.30E-17 | HNH endonuclease |
| ORF 3 | 1002 | reverse transcriptase/ maturase | RVT_N | 1.40E-29 | N-terminal domain of reverse transcriptase |
| | | | RVT_1 | 2.60E-26 | Reverse transcriptase (RNA-dependent DNA polymerase) |
| ORF 4 | 330 | HNH endonuclease | / | / | / |
| ORF 5 | 1809 | reverse transcriptase, intron maturase and HNH endonuclease | RVT_1 | 7.40E-46 | Reverse transcriptase (RNA-dependent DNA polymerase) |
| | | | RVT_N | 2.00E-25 | N-terminal domain of reverse transcriptase |
| | | | GIIM | 2.50E-19 | Group II intron, maturase-specific domain |
| ORF 6 | 2199 | reverse transcriptase, intron maturase | RVT_1 | 4.00E-23 | Reverse transcriptase (RNA-dependent DNA polymerase) |
| | | | Intron_maturas2 | 9.90E-27 | Type II intron maturase |
| ORF 7 | 2816 | reverse transcriptase, intron maturase | / | / | / |
| ORF 8 | 525 | HNH endonuclease | / | / | / |
| ORF 9 | 480 | HNH endonuclease | / | / | / |
| ORF 10 | 357 | HNH endonuclease | Intron_maturas2 | 7.00E-09 | Type II intron maturase |
| ORF 11 | 2448 | reverse transcriptase, intron maturase and HNH endonuclease | RVT_1 | 1.20E-21 | Reverse transcriptase (RNA-dependent DNA polymerase) |
| | | | Intron_maturas2 | 3.90E-21 | Type II intron maturase |
| ORF 12 | 582 | reverse transcriptase/ maturase | RVT_1 | 3.50E-13 | Reverse transcriptase (RNA-dependent DNA polymerase) |
| ORF 13 | 654 | LADILADG homing endonuclease | LAGLIDADG_2 | 1.90E-31 | LAGLIDADG DNA endonuclease family |
| ORF 14 | 1566 | | Asn_synthase | 6.90E-43 | Asparagine synthaseAsn_synthase |

| | | | | | |
|---|---|---|---|---|---|
| | | asparagine synthetase [glutamine-hydrolyzing] | GATase_7 | 1.70E-25 | Glutamine amidotransferase domain |
| ORF 15 | | LMBR1 domain containing protein | LMBR1 | 6.60E-32 | LMBR1-like membrane protein |
| ORF 16 | 687 | reverse transcriptase, intron maturase and HNH endonuclease | Intron_maturas2 | 1.30E-20 | Type II intron maturase |
| ORF 174 | 375 | LADILADG endonuclease | LAGLIDADG_1 | 1.40E-16 | LAGLIDADG endonuclease |
| ORF 18 | 456 | reverse transcriptase/ maturase | / | / | / |
| ORF 19 | 357 | reverse transcriptase/ maturase | GIIM | 3.30E-10 | Group II intron, maturase-specific domain |
| ORF 20 | 231 | reverse transcriptase/ maturase | RVT_1 | 8.40E-12 | Reverse transcriptase (RNA-dependent DNA polymerase) |
| ORF 21 | 252 | reverse transcriptase/ maturase | RVT_1 | 3.60E-14 | Reverse transcriptase (RNA-dependent DNA polymerase) |
| ORF 22 | 609 | reverse transcriptase/ maturase | RVT_1 | 2.50E-14 | Reverse transcriptase (RNA-dependent DNA polymerase) |
| ORF 23 | 162 | HNH endonuclease | / | / | / |
| ORF 24 | 696 | GIY-YIG homing endonuclease | / | / | / |
| ORF 25 | 210 | GIY-YIG homing endonuclease | / | / | / |
| ORF 26 | 507 | reverse transcriptase/ maturase | HNH | 4.30E-10 | HNH endonuclease |
| ORF 27 | 897 | reverse transcriptase, intron maturase and HNH endonuclease | / | / | / |
| ORF 28 | 495 | reverse transcriptase/ maturase | RVT_N | 1.10E-31 | N-terminal domain of reverse transcriptase |
| | | | RVT_1 | 6.50E-14 | Reverse transcriptase (RNA-dependent DNA polymerase) |
| ORF 29 | 981 | reverse transcriptase/ maturase | GIIM | 9.90E-20 | Group II intron, maturase-specific domain |
| | | | RVT_1 | 7.60E-14 | Reverse transcriptase (RNA-dependent DNA polymerase) |
| | | | HNH | 9.40E-12 | HNH endonuclease |
| ORF 30 | 2562 | reverse transcriptase, intron maturase and HNH endonuclease | RVT_1 | 1.60E-24 | Reverse transcriptase (RNA-dependent DNA polymerase) |
| | | | Intron_maturas2 | 7.20E-23 | Type II intron maturase |
| ORF 31 | 2376 | HNH endonuclease | RVT_1 | 2.50E-24 | Reverse transcriptase (RNA-dependent DNA polymerase)RVT_1 |
| | | | Intron_maturas2 | 1.90E-19 | Type II intron maturase |

| | | | | | |
|---|---|---|---|---|---|
| ORF 32 | 816 | HNH endonuclease | / | / | / |
| ORF 33 | 3396 | reverse transcriptase/ maturase | RVT_1<br>GIIM | 8.40E-20<br>3.70E-11 | Reverse transcriptase (RNA-dependent DNA polymerase)RVT_1<br>Group II intron, maturase-specific domain |
| ORF 34 | 636 | HNH endonuclease | / | / | / |
| ORF 35 | 1734 | HNH endonuclease | / | / | / |
| ORF 36 | 3576 | reverse transcriptase, intron maturase and HNH endonuclease | Intron_maturas2 | 9.10E-25 | Type II intron maturase |
| ORF 37 | 2121 | HNH endonuclease | RVT_1<br>Intron_maturas2 | 8.80E-15<br>3.10E-20 | Reverse transcriptase (RNA-dependent DNA polymerase)<br>Type II intron maturase |
| ORF 38 | 594 | GIY-YIG domain containing endonuclease | GIY-YIG | 1.10E-13 | GIY-YIG catalytic domain |
| ORF 39 | 555 | GIY-YIG domain containing endonuclease | GIY-YIG | 2.30E-11 | GIY-YIG catalytic domain |
| ORF 40 | 387 | reverse transcriptase/ maturase | RVT_N | 1.70E-27 | N-terminal domain of reverse transcriptase |
| ORF 41 | 531 | reverse transcriptase/ maturase | RVT_1 | 1.80E-23 | Reverse transcriptase (RNA-dependent DNA polymerase) |
| ORF 42 | 813 | HNH endonuclease | / | / | / |
| ORF 43 | 1782 | reverse transcriptase, intron maturase and HNH endonuclease | RVT_1<br>RVT_N<br>GIIM | 9.50E-37<br>2.00E-17<br>4.20E-16 | Reverse transcriptase (RNA-dependent DNA polymerase)<br>N-terminal domain of reverse transcriptase<br>Group II intron, maturase-specific domain |
| ORF 44 | 483 | reverse transcriptase | / | / | / |
| ORF 45 | 270 | reverse transcriptase/ maturase | RVT_1 | 1.20E-11 | Reverse transcriptase (RNA-dependent DNA polymerase) |
| ORF 96 | 963 | reverse transcriptase, II intron maturase / HNH endonuclease | RVT_1 | 3.80E-15 | Reverse transcriptase (RNA-dependent DNA polymerase) |
| ORF 165 | 627 | reverse transcriptase, II intron maturase | RVT_1 | 2.90E-13 | Reverse transcriptase (RNA-dependent DNA polymerase) |
| ORF 174 | 318 | group II reverse transcriptase | RVT_1 | 5.90E-18 | Reverse transcriptase (RNA-dependent DNA polymerase) |
| ORF 221 | 699 | reverse transcriptase, II intron maturase | / | / | / |

| | | | | | |
|---|---|---|---|---|---|
| ORF 230 | 1734 | reverse transcriptase, II intron maturase / HNH endonuclease | RVT_1 | 2.30E-12 | Reverse transcriptase (RNA-dependent DNA polymerase) |
| | | | intron_maturas2 | 1.50E-21 | Type II intron maturase |

**Table S7. Taxa considered in the RelTime analyses**

| Species | Strain or isolate; ptDNA GenBank accession |
|---|---|
| *Carteria crucifera* | UTEX 432; KT624880.1 |
| *Chlamydomonas applanata* | SAG 11-9; KT625417.1 |
| *Hyalomonas chlamydogama* | SAG 11-48b; Pending |
| *Chlamydomonas leiostraca* | SAG 11-49; NC_032109.1 |
| *Chlamydomonas reinhardtii* | CC-125; BK000554.2 |
| *Chlorella sorokiniana*[1] | Isolate 1230; KJ742376.1 |
| *Chlorella variabilis*[1] | NC64A; KJ718922.1 |
| *Chlorogonium capillatum* | UTEX11; KT625090.1 |
| *Chlorogonium euchlorum* | SAG 12-2a; NNNNN |
| *Chloromonas perforata* | SAG 11-43; KT625416.1 |
| *Chloromonas radiata* | UTEX 966; KT625083.1 |
| *Dunaliella salina* | CCAP 19/18; NC_016732.1 |
| *Gonium pectorale* | K3-F3-4 (NIES-2863); NC_020438.1 |
| *Haematococcus pluvialis* | UTEX 2505; NC_037007.1 |
| *Hyalogonium fusiforme* | SAG 62-1c; Pending |
| *Microglena monadina* | SAG 31.72; KT624802.1 |
| *Oltmannsiellopsis viridis* | NIES-360; NC_008099.1 |
| *Oogamochlamys gigantea* | SAG 44.91; NC_028580.1 |
| *Pectinodesmus pectinatus* | NA[2]; KU847995.1 |
| *Pleodorina starrii* | NIES-1363; NC_021109.1 |
| *Hyalomonas oviformis* | SAG 62-27: Pending |
| *Polytoma uvella* | UTEX 964; KX828177.1 |
| *Schizomeris leibleinii* | UTEX LB 1228; HQ700713.1 |
| *Stigeoclonium helveticum* | UTEX 441; DQ630521.1 |
| *Tetradesmus obliquus* | DOE0152 (UTEX B 3031); CM007919.1 |
| *Volvox carteri f. nagariensis* | UTEX 2908; GU084820.1 |

1. *Chlorella sorokiniana* and *Chlorella variabilis* were used as outgroup.
2. NA; Strain information is not available in the corresponding GenBank record.
Nonphotosynthetic species are indicated in shaded boxes.
